# Single-cell profiling reveals the impact of genetic alterations on the differentiation of inflammation-induced colon tumors

**DOI:** 10.1101/2023.11.30.569463

**Authors:** Ahmed H. Ghobashi, Rosie Lanzloth, Christopher A. Ladaika, Heather M. O’Hagan

## Abstract

Genetic mutations and chronic inflammation of the colon contribute to the development of colorectal cancer (CRC). Using a murine model of inflammation-induced colon tumorigenesis, we determined how genetic mutations alter colon tumor cell differentiation. Inflammation induced by enterotoxigenic *Bacteroides fragilis* (ETBF) colonization of multiple intestinal neoplasia (Min^ApcΔ716/+^) mice triggers loss of heterozygosity of *Apc* causing colon tumor formation. Here, we report that the addition of *BRAF*^V600E^ mutation (*BRAF*^FV600E^*Lgr5*^tm1(Cre/ERT2)Cle^Min^ApcΔ716/+^, BLM) or knocking out *Msh2* (*Msh2^LoxP/LoxP^Vil1-cre*Min^ApcΔ716/+^, MSH2KO) in the Min model altered colon tumor differentiation. Using single cell RNA-sequencing, we uncovered the differences between BLM, Min, and MSH2KO tumors at a single cell resolution. BLM tumors showed an increase in differentiated tumor epithelial cell lineages and a reduction in the stem cell population. In contrast, MSH2KO tumors were characterized by an increased stem cell population that had higher WNT signaling activity compared to Min tumors. Additionally, comparative analysis of single-cell transcriptomics revealed that BLM tumors had higher expression of transcription factors that drive differentiation, such as *Cdx2,* than Min tumors. Using RNA velocity, we were able to identify additional potential regulators of BLM tumor differentiation such as NDRG1. The role of CDX2 and NDRG1 as putative regulators for BLM tumor cell differentiation was verified using organoids derived from BLM tumors. Our results demonstrate the critical connections between genetic mutations and cell differentiation in inflammation-induced colon tumorigenesis. Understanding such roles will deepen our understanding of inflammation-associated colon cancer.

## Introduction

Colorectal cancer (CRC) is the third most common cause of cancer-related mortalities among males and females in the US (Siegel et al., 2023). Chronic inflammation is one of the major risk factors for CRC tumorigenesis (Chiba et al., 2012), as evidenced by patients with inflammatory bowel disease (IBD) having a higher risk of CRC development than individuals without IBD (Lucafò et al., 2021). Alterations in the gut microbiota can also contribute to intestinal inflammation and CRC (Fan et al., 2021). Enterotoxigenic *Bacteroides fragilis* (ETBF) is a strain of the common anaerobic gut bacteria, *Bacteroides fragilis,* that secretes a metalloprotease toxin and can lead to severe intestinal inflammation. ETBF is also associated with IBD development and increased risk of CRC incidence (Boleij et al., 2015; Fan et al., 2021; Ulger Toprak et al., 2006; Viljoen et al., 2015).

In addition to inflammation, genetic mutations play a key role in CRC initiation (Armaghany et al., 2012). The tumor suppressor gene Adenomatous Polyposis Coli *(APC)* is mutated in almost 85% of sporadic CRC (L. Zhang & Shay, 2017). ETBF colonization of multiple intestinal neoplasia (Min*^Apc^*^Δ716/+^) mice, which are heterozygous for mutant *Apc*, results in loss of heterozygosity (LOH) of the wildtype allele of *Apc* triggering colon tumor formation mainly in the distal part of the colon (Allen et al., 2022; Maiuri et al., 2017; Wu et al., 2009). Loss of the mismatch repair protein MSH2 causes microsatellite instability (MSI), which occurs in 10-20% of sporadic CRC (Armaghany et al., 2012). Hereditary loss of MSH2 is associated with Lynch syndrome, which increases the risk of different cancer development, including CRC (Armaghany et al., 2012; Li et al., 2021). We developed a mouse model in which mice have intestine-specific *Msh2* deletion driven by villin-cre and an *Apc* mutation (*Msh2^LoxP/LoxP^Vil1-cre*Min^ApcΔ716/+^, MSH2KO) (Maiuri et al., 2017). *BRAF*-activating mutations, which lead to the activation of the MAPK pathway, occur in almost 10% of CRC (Norreys et al., 1994). *BRAF* mutant CRC is characterized by poor overall survival and limited response to chemotherapies (Tabernero et al., 2022). To understand how *BRAF* mutation contributes to CRC development, we developed a mouse model in which *BRAF*^V600E^ expression is driven by Lgr5-cre in Min mice (*BRAF*^FV600E^*Lgr5*^tm1(Cre/ERT2)Cle^Min^ApcΔ716/+^, BLM) (Destefano Shields et al., 2021). We have previously demonstrated that loss of *Msh2* in Min mice increases distal colon tumorigenesis following ETBF colonization compared to Min mice (Maiuri et al., 2017). In contrast, ETBF colonization in BLM mice resulted in additional new tumors in the mid-proximal part of the colon (Destefano Shields et al., 2021), which resemble the right-sided location of *BRAF* mutant tumors observed in CRC patients (Tabernero et al., 2022). Furthermore, the BLM tumors exhibited a serrated and mucinous phenotype unlike Min tumors (Destefano Shields et al., 2021). Our previous works provide evidence for the role of gene mutation-inflammation interactions in inflammation-induced colon cancer tumorigenesis.

The intestinal colon epithelium is a continuously self-renewing tissue organized into defined crypt-villus units. Colon stem cells, located at the base of the crypt, undergo self-renewal and generate transit-amplifying cells, which migrate up the crypt-villus axis and differentiate into absorptive enterocytes and secretory cells (Clevers, 2013). The Wingless/Int (WNT)/β-catenin signaling pathway is essential to maintain stem cell proliferation at the bottom of the crypt (Clevers, 2013). Activating mutations of the WNT/β-catenin pathway, such as loss of *APC,* increases the stem cell population, which is associated with CRC initiation and progression (Schatoff et al., 2017). Genetic mutations and environmental factors, such as inflammation, disrupt the balance between stem and differentiated cells and contribute to CRC development.

This study aims to demonstrate the effect of gene mutation-inflammation interactions on colon tumor cell heterogeneity and differentiation. To accomplish this goal, we use single-cell RNA sequencing (scRNA-seq) of colon tumors collected from ETBF colonized Min, MSH2KO, and BLM mice to uncover cell type differences in the tumors at a single-cell resolution. We present here one of the first single-cell comparisons of inflammation-induced tumorigenesis in different genetic backgrounds. Our results demonstrate that genetic mutations induce tumor cell population differences and distinct stem cell differentiation patterns in inflammation-induced colon tumors. Determining how gene mutation-inflammation interactions alter the equilibrium between stem cells and differentiated cells will enhance our understanding of inflammation-associated colon tumorigenesis.

## Results

### Single-cell profiling identifies cell populations in colon tumors

To investigate if *BRAF* mutation or microsatellite instability caused by *Msh2* deletion modifies the cellular composition of ETBF-driven colon tumors, ETBF colonization of BLM, MSH2KO, and Min C57BL/6 mice was performed. For each sample, colon tumors were harvested after 6-8 weeks of ETBF colonization and enzymatically digested. Then, scRNA-seq was performed using droplet-based microfluidics (10X Genomics) (Fig. 1A). scRNA-seq of tumors from BLM mice from two different experimental cohorts was performed with the samples labeled BLM and BLM2. The number of total single cells initially sequenced from each sample was BLM:12,273, BLM2:12,733, MSH2KO:9,618, and Min:7,535.

**Figure 1.**
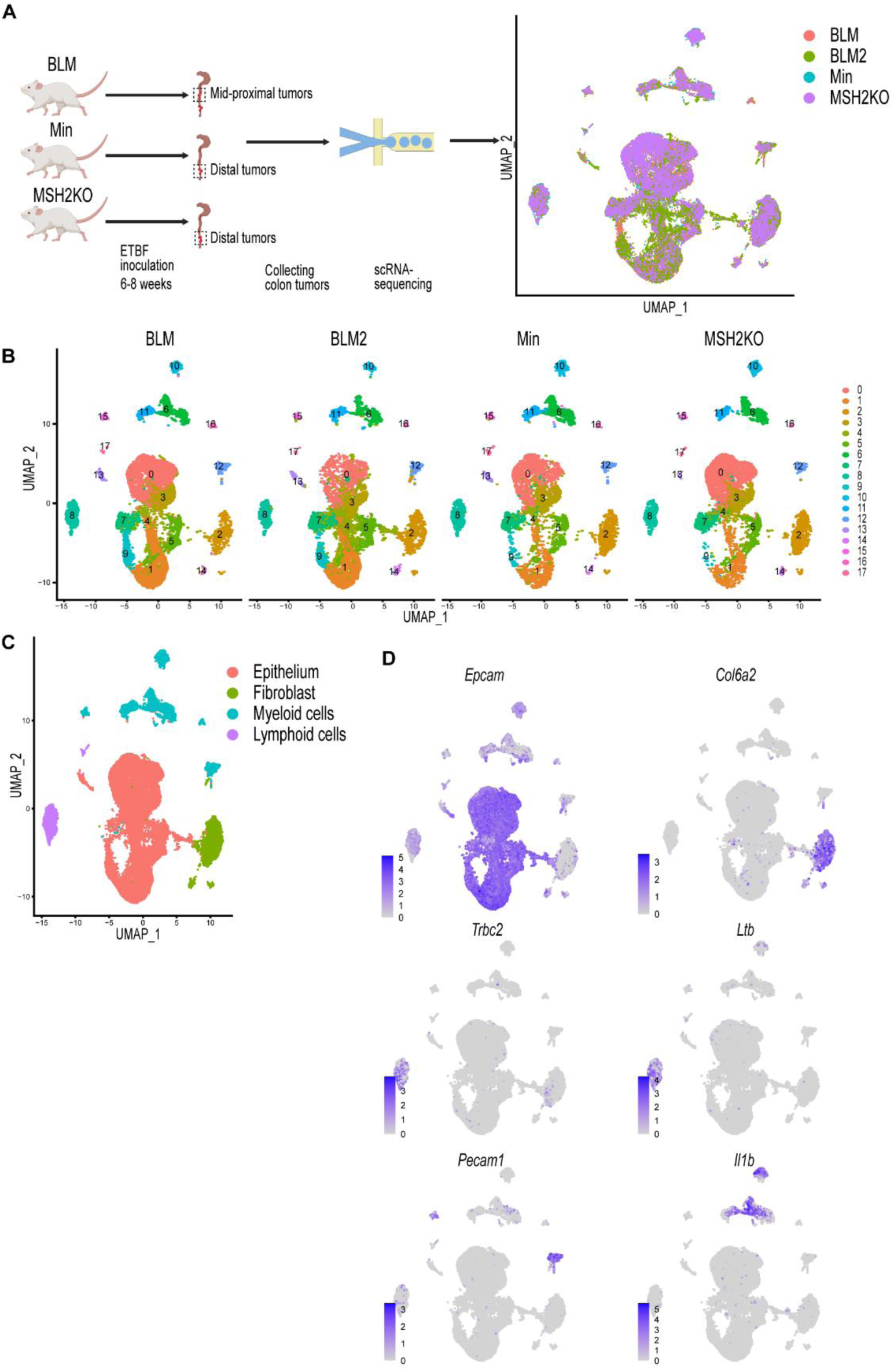
Single cell profiling identifies cell populations in colon tumors. (A) Diagrammatic representation of sample preparation and single cell RNA-sequencing (scRNA-seq) and uniform manifold approximation and projection (UMAP) plot of the four integrated samples. (B) UMAP plot of BLM, BLM2, Min, and MSH2KO colon tumor scRNA-seq samples colored by cluster. (C) UMAP plot of the four integrated scRNA-seq samples colored by major cell type. (D) Featureplot of normalized expression values of marker genes representative of the epithelial, fibroblast, lymphoid, and myeloid cell populations in the combined scRNA-seq samples. Color intensity represent the normalized gene expression.

Quality control was performed by removing cells with mitochondrial content higher than 20% and removing cell doublets. After cell removal, the number of cells for each sample was BLM:10,062, BLM2:9,946, MSH2KO:8,030, and Min:6,337. Using Seurat (Hao et al., 2021), data were normalized using sctransform, and 3,000 input variable genes were used to identify integration anchors among the four different datasets. Following integration, dimensional reduction, and unsupervised Louvain modularity-based clustering were performed resulting in 18 clusters (Fig. 1B). Visualizing these subpopulations using uniform manifold approximation and projection (UMAP, Fig. 1B) confirmed their distinct identities. To identify different major cell populations (epithelial cells, immune cells, and fibroblasts), we manually assigned class identities based on the expression of well-established marker genes (Fig. 1C). Epithelial cells were identified through the expression of *Epcam, Krt8,* and *Krt18* (Fig. 1D, Supplemental Fig. S1). *Col6a2* and *Pdgfrb* were used to identify fibroblasts (Fig. 1D, Supplemental Fig. S1). *Trbc2, Ltb,* and Emb were used to annotate lymphoid cells, and *Pecam1*, *Il1b Cd68*, and *Lyz2* were used to identify myeloid cells (Fig. 1D, Supplemental Fig. S1).

### Single cell survey of colon tumor epithelial cells

To focus our study on colon tumor differentiation, we extracted the epithelial cells from the dataset and reclustered them to produce 10 distinct epithelial cell clusters (Fig. 2A). Proliferating cells were identified using cell-cycle signatures (Fig. 2B). Seurat FindAllMarkers was utilized to identify transcripts enriched for each cluster, and then each cluster was manually annotated (Supplemental Table S1). We identified a stem cell cluster (cluster 0) with *Axin2*, *Sox4*, *Ccnd2*, and *Clu* expression (Fig. 2C, Supplemental Fig. S2). Transit-amplifying cells (TA; cluster 2) were enriched for proliferating markers; *Ccna2*, *Pcna*, *Hmgn2*, *Mcm5*, *Top2a*, *Mki67* (Fig. 2C, Supplemental Fig. S2), which is consistent with cell-cycle signature result (Fig. 2B). Paneth cells (PC; cluster 7) were identified by the expression of the anti-microbial genes *Lyz1*, *Ang4*, *Spink4*, and *Mmp7* (Fig. 2C, Supplemental Fig. S2). Goblet cells (GC; cluster 5) were marked by the elevated expression of *Zg16*, *Fcgbp*, *Tff3*, *Muc2*, and *Sval1* (Fig. 2C, Supplemental Fig. S2). Cluster 1 was labeled as enterocyte cells (EC) due to its enrichment with *Car1, Fabp2*, *Slc26a3*, and *Ndrg1* (Fig. 2C, Supplemental Fig. S2). Enterocytes/brush-border cells (cluster 4) were marked by elevated levels of *Cdhr5, Car4*, *Aqp8*, *Guca2a*, *Saa1*, and *Espn* (Fig. 2C, Supplemental Fig. S2). Cluster 8 was characterized by the expression of markers that are normally expressed in fibroblasts such as *Saa3*, *Dcn, Lgals1, and Bmp4* (Fig. 2C, Supplemental Fig. S2). Because several of these genes, such as *Saa3* and *Dcn*, have previously been shown to be expressed in goblet cells (Gorman et al., 2023; Reigstad & Bäckhed, 2010), this cluster was labeled “secretory-like cells” (Fig. 2C, Supplemental Fig. S2). Additionally, to confirm that this cluster is indeed part of the colon tumor epithelium, we performed IHC for SAA3. While we detected SAA3-positive cells in the epithelial tumor tissue, Saa3-stained cells were mainly stromal cells in the normal colon (Fig. 2D). To follow up on our IHC result, we performed immunofluorescence staining for both SAA3 and E-cadherin (an epithelial cell marker). In BLM tumor tissue, unlike normal colon tissue, E-cadherin-positive cells were also positive for SAA3 protein (Fig. 2E) further confirming that the *Saa3*-expressing cluster (secretory-like cells) is part of the colon tumor epithelium. Interestingly, SAA3 appeared to be nuclear in the tumor cells but cytoplasmic in the stromal cells in the normal colon (Fig. 2D, 2E). Cluster 9 was removed because the cells were enriched for genes predominantly expressed in immune cells, such as *Lyz2*, *S100a9*, *Il1b*, and *S100a8* (cluster 9, Supplemental Table S1). Clusters 3 and 6 were also excluded because they expressed *AY036118* and *Gm26917,* which are associated with ribosomal RNA contamination (Supplemental Table S1).

**Figure 2.**
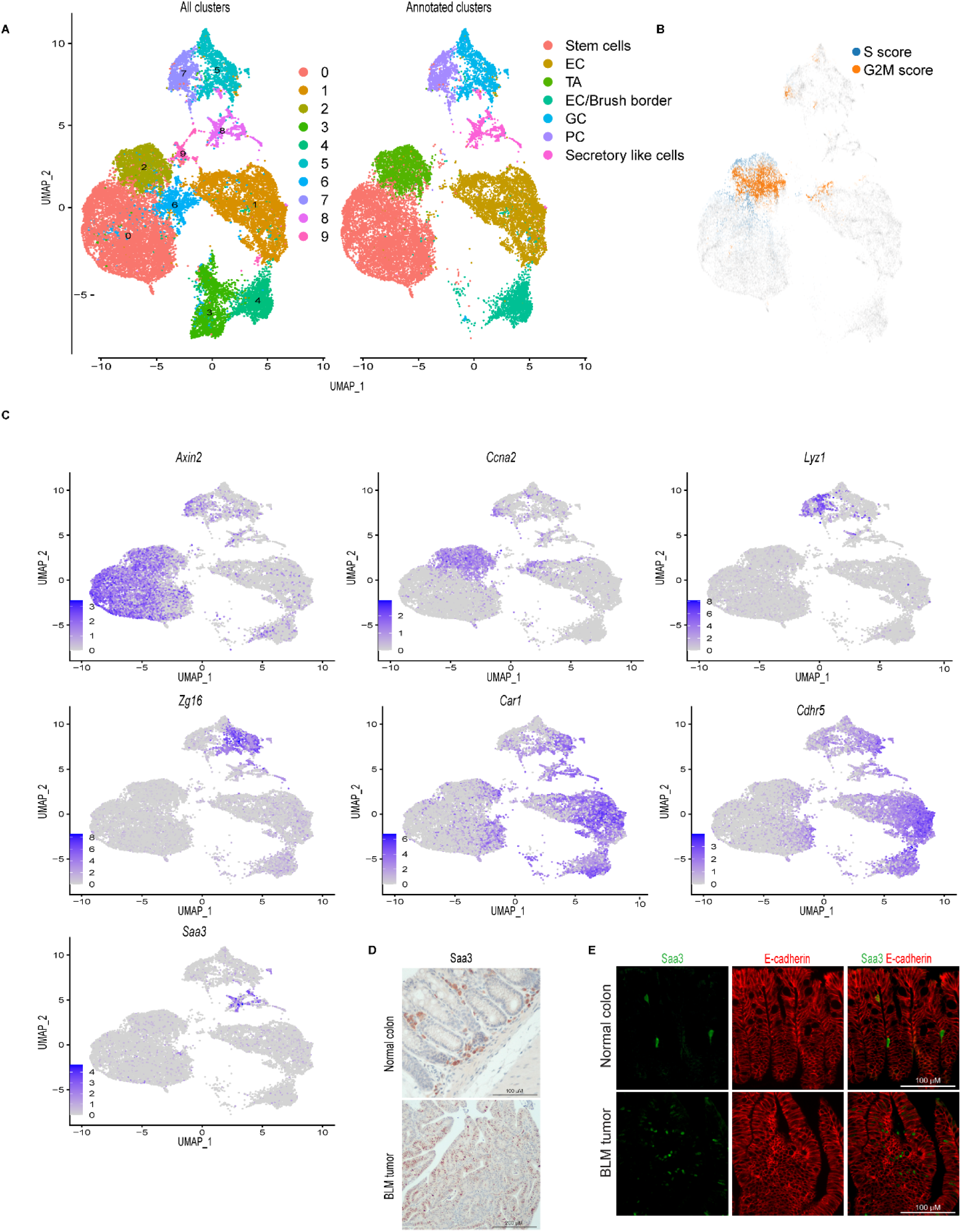
Single cell survey of colon tumor epithelial cells. (A) Uniform manifold approximation and projection (UMAP) plot of the combined colon epithelial tumor scRNA-seq data colored by cluster (left, All clusters) and by cell type after filtering out poor quality clusters (right, Annotated clusters; enterocyte cells (EC), transit-amplifying (TA), goblet cells (GC), Paneth cells (PC)). (B) UMAP plot of the combined scRNA-seq samples showing the expression of cell cycle signature genes (S score and G2M score). (C) Featureplots of normalized expression values of marker genes for the different clusters/cell types. Color intensity represent the normalized gene expression. (D) Representative SAA3 IHC of BLM normal colon (Scale bar, 100 μm) and BLM tumor (Scale bar, 200 μm). (E) Representative SAA3 (green) and E-cadherin (red) immunofluorescent images of BLM tumor and BLM normal colon (Scale bar, 100 μm).

### *BRAF^V600E^* mutation and *Msh2* deletion alter colon tumor epithelial cell composition

To investigate the effect of the expression of the *BRAF^V600E^*mutation or *Msh2* deletion on epithelial cell composition in inflammation-induced colon tumors, we compared the proportion of cells in the different colon tumor cell populations between the different tumor types using Min tumors as a control (Fig. 3A, 3B). Interestingly, *BRAF* mutant tumors (BLM and BLM2) were enriched for differentiated cell populations such as secretory-like cells, goblet cells, and EC/brush border cells compared to Min tumors (Fig. 3B, BLM vs Min and BLM2 vs Min). In contrast, *BRAF* mutant tumors had significantly reduced stem cell and TA cell populations compared to Min tumors (Fig. 3B, BLM2 vs Min). These results agree with other findings suggesting that *BRAF* mutation induces colon epithelial cell differentiation (Tong et al., 2017). While *Msh2* deleted tumors had a significantly increased stem cell population, they had reduced differentiated cell populations such as secretory-like cells, enterocytes, and goblet cells in comparison to Min tumors (Fig. 3B, MSH2KO vs Min). Next, we compared the expression of different transcripts that are known to be specific to each lineage. In agreement with more differentiated cells in BLM tumors, BLM tumors had higher expression of *Muc2* (Goblet cells), *Fabp2* (enterocytes), and *Saa3* (secretory-like cells) than Min and MSH2KO tumors (Supplemental Fig. S3). Consistent with the significant increase of secretory-like cells in BLM2 (Fig. 3B, BLM2 vs Min), tumor cells in BLM tumors showed a significant increase in SAA3 protein levels compared to Min tumors by IHC (Fig. 3C, 3D). Paneth cell marker, *Lyz1*, was expressed more in Min and MSH2KO tumor paneth cells than BLM tumors (Supplemental Fig. S3), which is consistent with the smaller Paneth cell population in BLM tumors (Fig. 3B, BLM vs. Min). MSH2KO and Min tumors also had a higher percentage of cells in the TA and stem cell clusters expressing *Sox4* than BLM tumors (Supplemental Fig. S3). Interestingly, while *Reg3g* and *Guca2a* were primarily restricted to Paneth cells in Min and MSH2KO tumors, they were also expressed in enterocytes of BLM tumors (Supplemental Fig. S3). Because *Guca2a* was expressed in multiple lineages, we computed RNA velocity for *Guca2a* to calculate its differential RNA splicing kinetics allowing us to determine which lineages display RNA splicing kinetics for the *Guca2a* transcript in the different tumor types (see methods). We found that in BLM tumors, *Guca2a* exhibited high velocity (high unspliced to spliced RNA ratio) in goblet and enterocyte cells with significant differential RNA splicing kinetics in enterocytes (Fit pval kinetics= 2.29E-07) (Fig. 3E, BLM2). However, in Min tumors, Paneth cells and TA showed significant differential kinetics for the *Guca2a* transcript (Fit pval kinetics= 5.99E-06) (Fig. 3E, Min).

**Figure 3.**
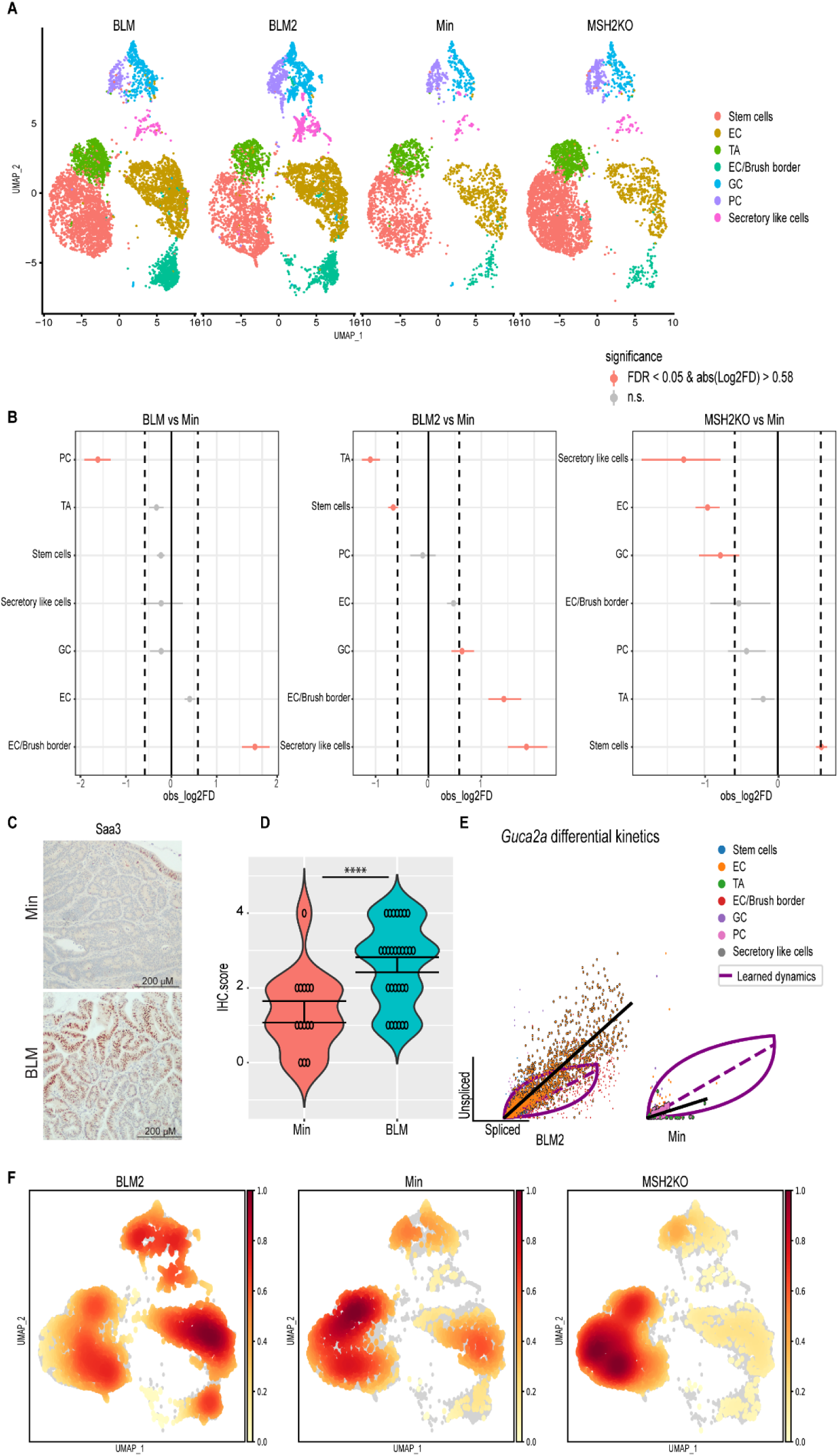
*BRAF* mutation and *Msh2* deletion have different tumor epithelial cellular compositions. (A) Uniform manifold approximation and projection (UMAP) plot of the BLM, BLM2, Min, and MSH2KO colon tumor epithelial cells colored by cell type (enterocyte cells (EC), transit-amplifying (TA), goblet cells (GC), Paneth cells (PC)). (B) Relative differences in cell proportions for each cluster between the BLM versus Min, BLM2 versus Min, and MSH2KO versus Min samples. Red dots have an FDR < 0.05 and mean |Log 2-fold difference (Log2FD)|>0.58 compared with the Min colon tumor (permutation test; n = 10,000). (C) Representative SAA3 IHC in BLM and Min colon tumors (Scale bar, 200 μm). (D) Quantification of SAA3 IHC stain in (C). (E) Scatter plot for the unspliced and spliced transcript for *Guca2a* calculated by RNA velocity. The black line represents the significant differential RNA splicing kinetics for enterocytes (BLM2) and Paneth cells (Min) and the purple line represents the overall RNA dynamic. Cells are colored by cell type. (F) Embedding density plot of BLM2, Min, and MSH2KO samples. Colors represent the scaled density values. Significance was determined by paired t-test. **** p <= 0.0001.

To confirm that BLM tumors were more differentiated than Min tumors, we calculated the density of cells in 2D space. Consistent with our findings, BLM tumors showed more cell density toward the differentiated enterocyte lineages, while the TA cluster had more cell density in Min tumors (Fig. 3F). There was also more cell density in the stem cell population in MSH2KO tumors compared to Min and BLM tumors (Fig. 3F, MSH2KO). Our findings suggest that expression of mutant *BRAF^V600E^* or deletion of *Msh2* alters cell composition in inflammation-induced colon tumors with increased and decreased differentiated cells in BLM and MSH2KO tumors, respectively.

### *BRAF^V600E^* colon tumors are more differentiated than Min and *Msh2* deleted tumors

We hypothesized that the difference in tumor cellular composition among the different datasets is due to differences in colon tumor cell differentiation. To better understand colon tumor epithelial cell differentiation in the different tumor types, computational trajectory analysis was performed. For all samples, a cyclical pattern emerged representing cycling stem cells and TA cells that also branched to secretory lineages and enterocytes (Fig. 4A). While the BLM tumors had more cells moving toward differentiated lineages (Fig. 4A, BLM2), MSH2KO tumors appeared to have more cycling stem cells in comparison to Min tumors (Fig. 4A, MSH2KO). Additionally, we computed velocity confidence and velocity length to predict cell directionality and differentiation speed, respectively. Compared to Min tumors, BLM tumors showed more directionality toward differentiated lineages such as goblet cells, enterocytes, and brush border cells with an increased rate of differentiation at enterocytes and brush border cells (Fig. 4B, BLM2). MSH2KO tumors showed more directionality and speed toward the stem cell population compared to Min tumors (Fig. 4B, MSH2KO).

**Figure 4.**
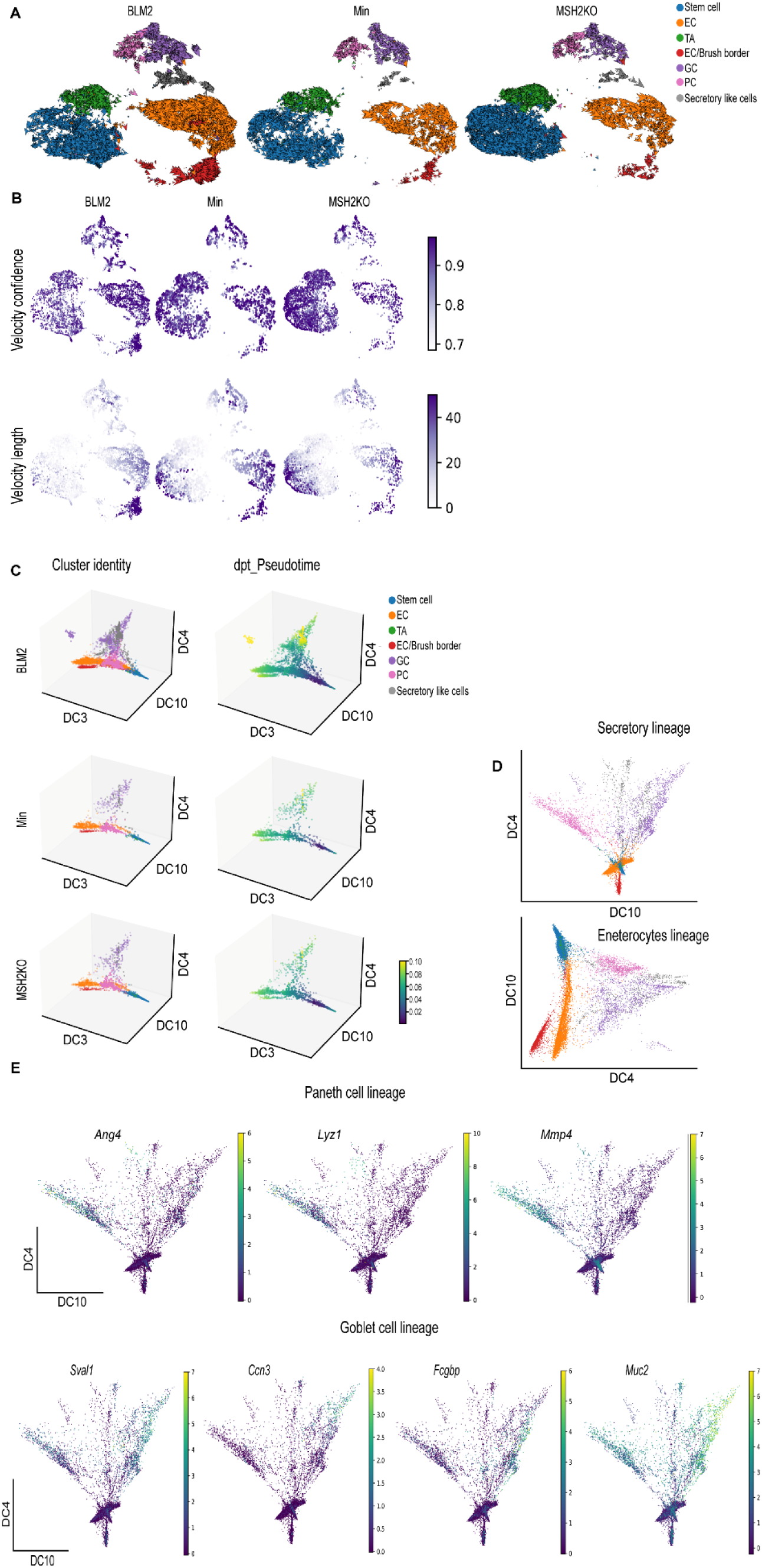
*BRAF* mutant colon tumors are more differentiated than Min and *Msh2* deleted tumors. (A) RNA velocity arrows for individual cells of the BLM2, Min, and MSH2KO tumor epithelial scRNA-seq data colored by cell type, showing inferred differentiation trajectories (enterocyte cells (EC), transit-amplifying (TA), goblet cells (GC), Paneth cells (PC)). (B) Uniform manifold approximation and projection (UMAP) plot showing RNA velocity confidence and length in BLM2, Min, and MSH2KO samples. Color intensity represents the velocity confidence and length values. (C) The diffusion-map embeddings of BLM2, Min, and MSH2KO colon tumor epithelial cells are colored by cell type (left, Cluster identity) and diffusion pseudotime (right, dpt_Pseudotime). Diffusion components (DCs) 3, 4, and 10 correspond approximately to the colon tumor differentiation state. Colors represent pesudotime values. (D) Diffusion-map embedding of combined BLM, BLM2, Min, and MSH2KO colon tumor epithelium. DCs 4 and 10 capture secretory lineage differentiation (top) and DCs 3 and 4 capture enterocyte lineage differentiation (bottom). (E) Expression of regional markers of Paneth cell lineage (top) and goblet cell lineage (bottom) in the combined BLM, BLM2, Min, and MSH2KO tumor epithelial scRNA-seq data. Colors represent the normalized gene expression.

We next used diffusion maps to place the colon tumor epithelial populations in pseudo-temporal order (Haghverdi et al., 2016) (Fig. 4C, Supplemental Fig. S4A). We observed a trajectory from stem cells to the different differentiated lineages (Fig. 4C, cluster identity) and captured distinct paths towards enterocytes, goblet cells, secretory-like cells, and paneth cells (Fig. 4C, cluster identity). Importantly, the secretory-like cells were on the same trajectory as the other secretory cell types (goblet and Paneth cells), further supporting our identification of them as a secretory-like population. We also computed Diffusion Pseudotime (dpt) to calculate cell progression during differentiation. Consistent with more epithelial cell differentiation in BLM tumors, we observed more differentiated cells (yellow) on the pseudotime scale in BLM tumors compared to Min tumors (Fig. 4C, dpt_pseudotime). Using diffusion components 4 and 10 only, we were able to get better separation of the different secretory lineages (Fig. 4D, secretory lineages). Improved trajectory separation for enterocytes/brush border cells was accomplished using diffusion components 3 and 4 (Fig. 4D, enterocyte lineage). By identifying transcripts that were expressed in different regions of the diffusion map, we were able to associate the expression of genes with cell fate commitment to Paneth cells (*Ang4*, Lyz1, *Mmp7*; Fig. 4E), goblet cells (*Sval1*, *Ccn3, Fcgbp*, *Muc2*; Fig. 4E), secretory-like cells (*Saa3*, *Dcn, Bmp4*; Supplemental Fig. S4B), and enterocytes and brush border cells (*Saa1, Aqp8, Cdhr5*, *Muc3*, *Mall*; Supplemental Fig. S4B).

### Stem cells of MSH2-deleted tumors have high WNT signaling activity

In the normal intestine, the activity of the WNT/β-catenin signaling pathway is highest in the intestinal stem cells at the bottom of the crypts, where it regulates stem cell proliferation and differentiation (Clevers, 2013). Because MSH2KO and BLM tumors showed high cell density at stem cell and enterocyte clusters, respectively, we hypothesized that WNT activity would differ between tumor types and cell clusters even though all tumors are from mice on the Min background. Intriguingly, while stem cells of MSH2KO tumors showed high levels of WNT signaling pathway activity compared to Min tumors, stem cells in BLM tumors showed low WNT signaling activity (Fig. 5A). Next, we calculated differential gene expression between the stem cell clusters in MSH2KO and Min tumors. MSH2KO tumor stem cells had increased expression of WNT and stemness-related genes such as *Axin2*, *Tcf4*, *Wnt6*, *Wnt10a*, and *Prox1* relative to Min tumor stem cells (Fig. 5B). *Wnt6* and *Wnt10a* were highly expressed in stem cell populations of MSH2KO tumors compared to Min tumors and were almost absent in BLM tumor stem cells (Supplemental Fig. S5A). Additionally, organoids derived from MSH2KO tumors had significantly increased *Axin2* expression compared to organoids derived from Min tumors (Fig. 5C). Gene ontology (GO) analysis demonstrated that upregulated genes in MSH2KO stem cells were enriched for the WNT signaling pathway (Supplemental Fig. S5B). Consistent with the role of WNT signaling in maintaining the stem cell population and BLM tumors having fewer stem cells, BLM tumor stem cells had decreased expression of WNT/stemness-related genes compared to Min tumors such as *Axin2*, *Id2, Sox4*, *Wnt6*, *Wnt10a*, *Id3*, *Prox1*, and *Id1* (Fig. 5D, Supplemental Fig. S5C). In contrast, the stem cell population of BLM tumors had significant up-regulation of differentiation-related genes such as *Guca2a, Car1, Cdx2*, *Car2*, and *Muc2* (Fig. 5D, Supplemental Fig. S5D). Additionally, organoids derived from BLM2 tumors had a significant reduction in the expression of *Lgr5*, *Axin2*, and *Sox4* compared to Min organoids (Fig. 5E). Furthermore, up-regulated genes in BLM tumor stem cells showed significant enrichment for epithelial cell differentiation and regulation of microvillus length by GO analysis (Supplemental Fig. S5D). To confirm that the reduction of cycling stem cells in BLM2 is due to the reduced expression of stemness-related genes, we simulated the over-expression (OE) of *Id2,* a gene that was mainly expressed in the stem cells of BLM2 tumors (Fig. 5F) and was significantly down-regulated in stem cells of BLM tumors compared to Min tumors (Fig. 5D, Supplemental Fig. S5C). Interestingly, simulated OE of *Id2* in BLM2 caused a shift in TA and differentiated cells toward the stem cell population with more cycling TA and stem cells (Fig. 5F compared to BLM2 in Fig. 4A). These findings suggest that increased WNT signaling activity in the stem cell population of MSH2KO tumors contributes to the increased number of stem cells and reduced differentiation in these tumors.

**Figure 5.**
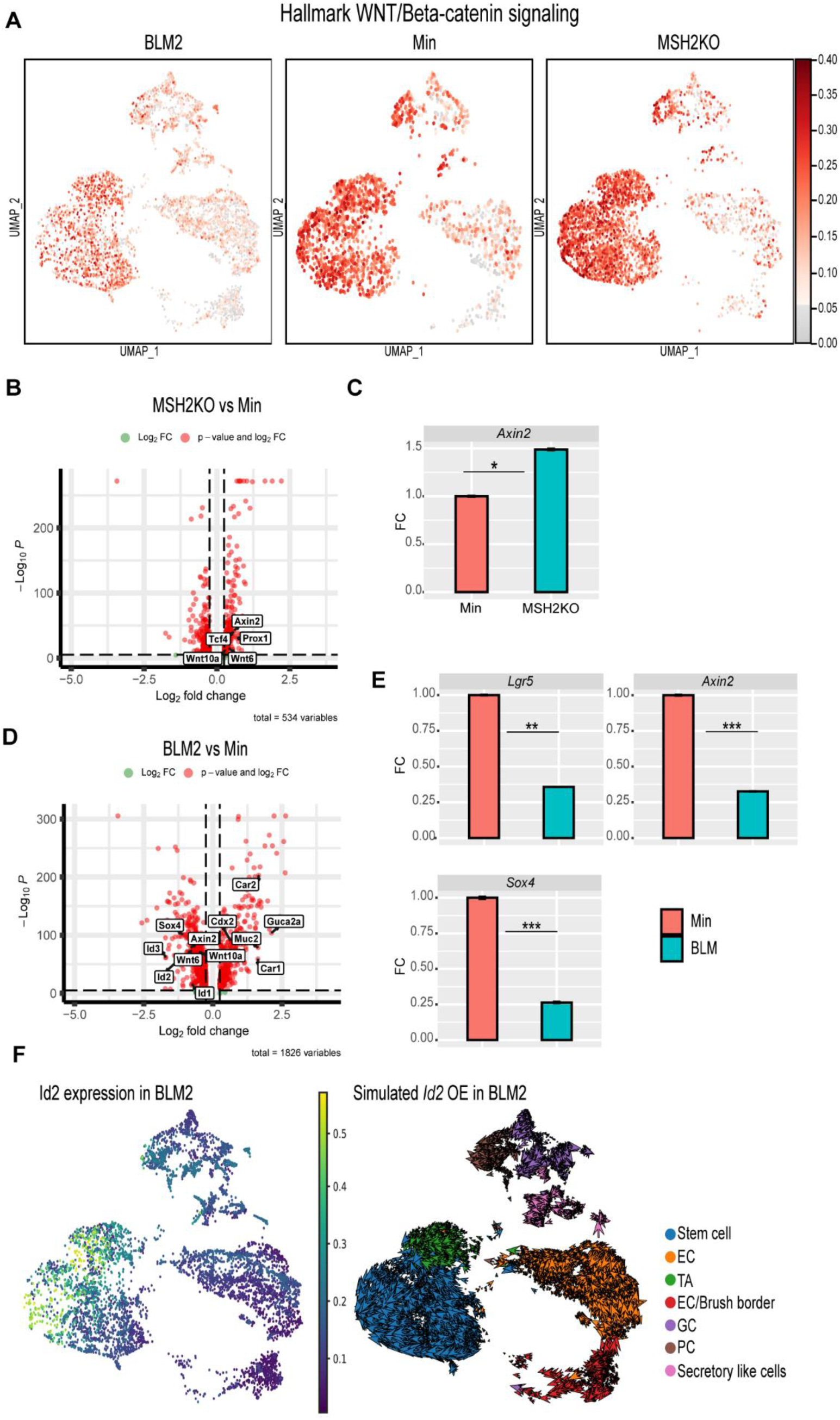
Stem cells of BLM and MSH2KO colon tumors have low and high WNT signaling activity, respectively. (A) Uniform manifold approximation and projection (UMAP) plot of BLM2, Min, and MSH2KO tumor epithelial scRNA-seq data showing the Hallmark WNT/β-catenin signaling score in each cell. Color intensity represents WNT signaling score (B) Volcanoplot of differentially expressed genes (DEGs) in the stem cell populations of MSH2KO versus Min. Dashed lines indicate |log 2FC|>0.25 and P<0.05. (C) Gene expression of the indicated genes by RT-qPCR. Expression of all the genes was normalized to the housekeeping gene *Ppia* and then to Min organoids. Results are represented as the mean of 3 biological replicates +/− SEM. (D) Volcanoplot of DEGs in the stem cell populations of BLM2 versus Min. Dashed lines indicate |log 2FC|>0.25 and P<0.05. (E) Gene expression of the indicated genes by RT-qPCR as in C. (F) UMAP plot showing the expression of *Id2* in BLM2 tumor epithelial cells (left) and RNA velocity arrow for individual cells in simulated *Id2* overexpression in BLM2 colon tumor (right). Colors represent the normalized gene expression. Significance was determined by paired t-test. *p <= 0.05, **p <= 0.01, ***p <= 0.001.

### CDX2 is involved in *BRAF^V600E^* colon tumor epithelial cell differentiation

So far, our data suggest that BLM tumors are more differentiated than Min tumors. To identify additional genes whose velocity drives toward different differentiation trajectories, cluster-specific differential velocity expressions were computed. For the secretory lineage, individual genes such as *Spdef* were identified, which are known to regulate secretory cell (Paneth and goblet cell) specification in the normal intestine (Lo et al., 2017), as well as additional novel regulators such as *Sytl2* and *Pld1* (Fig. 6A, Secretory lineage, Supplemental Table S2). Genes were identified that are associated with Paneth cell differentiation trajectories such as *Kcnb2*, *Hepacam2*, *Foxa3*, *Ern2*, and *Klf7* suggesting these genes may be novel regulators of Paneth cell differentiation (Fig. 6A, Paneth lineage, Supplemental Table S2). *Muc2* and *Stim1*, which are known to regulate goblet cell differentiation (Chang et al., 1994; Liang et al., 2022), as well as other novel goblet cell regulators such as *Muc4*, *Dstn*, and *Muc13,* were identified (Fig. 6A, Goblet lineage, Supplemental Table S2). Secretory-like cell regulators were identified such as *Nrg1*, *Piezo2,* and *Ncam1* (Fig. 6A, Secretory-like cell lineage, Supplemental Table S2). *Parm1*, *Mpp5, Muc3*, *Syk*, *Ascl3*, *Prag1*, *Cdh17*, and *Sgk2* were identified as potential regulators for enterocyte cell and enterocyte/brush border cell specification (Fig. 6A, Enterocytes and brush border lineage, Supplementary Table S2).

**Figure 6.**
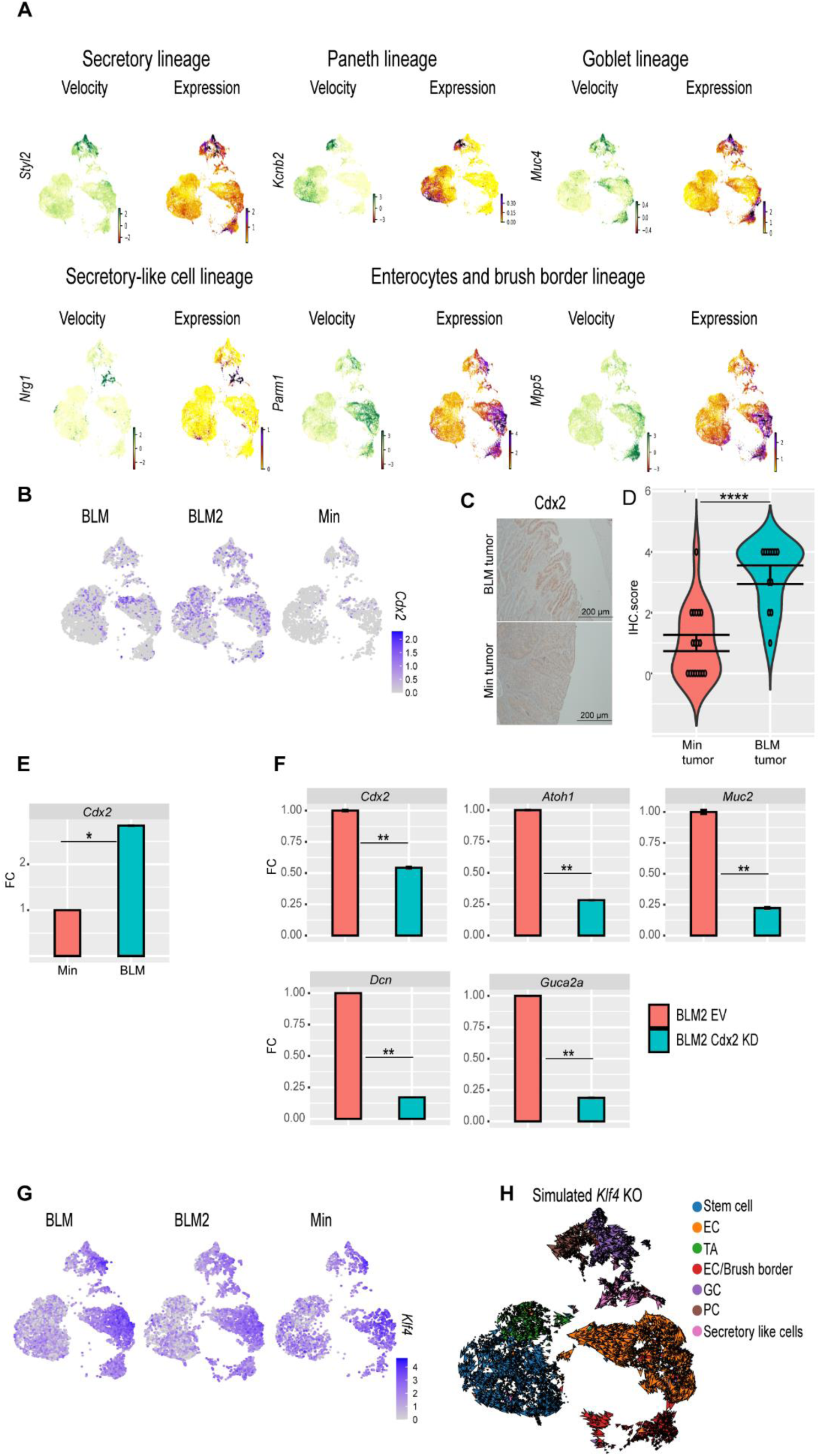
scRNA-seq reveals genes involved in BLM colon tumor epithelial cell differentiation. (A) Velocity and expression Uniform manifold approximation and projection (UMAP) plots of indicated genes that have differential velocity in the indicated cell lineages in combined tumor epithelial scRNA-seq data. Colors represent normalized gene and velocity expression. (B) Featureplot of normalized *Cdx2* expression in BLM, BLM2, and Min colon tumor epithelial cells. Color intensity represents normalized *Cdx2* gene expression. (C) Representative CDX2 IHC in BLM and Min colon tumors (Scale bar, 200 μm). (D) Quantification of Cdx2 IHC in (C). (E) and (F) Gene expression of the indicated genes in tumor organoids by RT-qPCR. Expression of all the genes was normalized to the housekeeping gene *Ppia* and then to Min organoids in (E) or to empty vector (EV) BLM2 organoids in (F). Results are represented as the mean of 3 biological replicates +/− SEM. (G) Featureplot of normalized *Klf4* expression in BLM, BLM2, and Min colon tumor epithelium samples. (H) UMAP plot showing the RNA velocity arrows for individual cells in simulated *KLF4* knockout in BLM2 colon tumor epithelial cells. Significance was determined by paired t-test. *p <= 0.05, **p <= 0.01, **** p <= 0.0001.

The CDX2 transcription factor plays an essential role in the development of the intestinal epithelium (Gendron et al., 2006; Kumar et al., 2019). Because stem cells of BLM tumors showed increased expression of *Cdx2* (Fig. 5D), we focused on CDX2 as a potential regulator of BLM tumor differentiation. BLM tumors had more *Cdx2*-expressing cells than Min tumors (Fig. 6B). Furthermore, by IHC, BLM tumor tissue had higher levels of CDX2 protein than Min tumors (Fig. 6C, 6D), whereas normal Min and BLM colon tissue had similar levels of CDX2 protein (Supplemental Fig. S6A). These findings suggest that inflammation-induced colon tumorigenesis induces loss of *Cdx2* expression in Min mice. Additionally, BLM organoids had significantly higher expression of *Cdx2* than Min organoids (Fig. 6E). To test the effect of CDX2 loss on differentiation, we knocked down *Cdx2* in organoids derived from BLM and BLM2 tumors (Fig. 6E, Supplemental Fig. S6B). Interestingly, knocking down *Cdx2* significantly reduced the expression of *Atoh1* (secretory cells), *Muc2* (Goblet cells), *Dcn* (Secretory-like cells), and *Guca2a* (Goblet and enterocyte cells) (Fig. 6F, Supplemental Fig. S6B). These results suggest that the expression of *Cdx2* in BLM tumor cells contributes to their differentiation.

In addition to CDX2, KLF4 is another transcription factor that plays an essential role in colon epithelium differentiation (Yu et al., 2012). *Klf4* had the highest levels of expression in differentiated cells, including goblet, secretory-like cells, and enterocyte cells (Fig. 6G). To test if disruption of *Klf4* expression affects BLM tumor differentiation, we simulated *Klf4* knockout (KO). Simulated *Klf4* KO changed the directionality of differentiated cells of BLM2 tumors, shifting their direction toward the stem cells (Fig. 6H, compared to BLM2 in Fig. 4A). This finding suggests that *Klf4* expression in BLM tumors may also contribute to BLM epithelial tumor differentiation.

To identify other potential driver genes for BLM tumor differentiation, we computed transcriptional dynamics using RNA velocity, which identified *Ndrg1* as a potential regulator of BLM tumor epithelial cell differentiation. NDRG1 has previously been shown to be involved in the differentiation of adipocytes and macrophages (Cai et al., 2017; Watari et al., 2016). The enterocyte clusters had the highest level of *Ndrg1* expression (Figure S6C, Expression). *Ndrg1* also had high velocity in enterocytes, and almost all the enterocyte populations showed high *Ndrg1* velocity in BLM and Min tumors (Figure S6C, Velocity). Interestingly, *Ndrg1* had high velocity in the stem cells of BLM tumors, while in Min tumors, *Ndrg1* only started to show velocity in differentiated enterocytes and goblet cells (Figure S6C, Velocity). This observation suggests that the high velocity of *Ndrg1* in stem cells might cause stem cells to differentiate in BLM tumors. Additionally, knocking down *Ndrg1* in BLM tumor organoids reduced secretory progenitor marker gene expression (Figure S6D; *Sox4*, *Atoh1*, *Dll1*, *Notch1*), but increased enterocyte marker gene expression (Figure S6D; *Cdhr2*, *Aqp8*). These results suggest that Ndrg1 may be an additional regulator of BLM tumor epithelial cell differentiation.

## Discussion

Recent advances in scRNA-seq technology have allowed for the evaluation of the intestinal epithelium, providing insights into the complexity and diversity of intestinal epithelium cell populations (Haber et al., 2017; Hickey et al., 2023). However, limited studies have examined the role of gene mutation and inflammation on cellular heterogeneity of intestinal epithelial tumors at a single-cell resolution. We previously demonstrated the impact of genetic mutations on phenotypic and molecular characteristics of inflammation-induced colon tumorigenesis (Destefano Shields et al., 2021; Maiuri et al., 2017). Many studies have demonstrated that genetic mutations and tumor heterogeneity play an enormous role in the effectiveness of chemotherapeutic treatments (Dagogo-Jack & Shaw, 2018; Waarts et al., 2022). Therefore, utilizing scRNA-seq, we focused on evaluating the changes in the cellular composition and differentiation of inflammation-induced murine colon tumors from three different genetic backgrounds (Min, BLM, and MSH2KO). We used ETBF colonization to induce colon inflammation because ETBF colonization of colon mucosa is associated with CRC incidence in humans (Boleij et al., 2015; Fan et al., 2021; Ulger Toprak et al., 2006; Viljoen et al., 2015). Importantly, ETBF induces LOH of *Apc* to trigger colon tumor formation without causing additional genetic mutations making it ideal to study inflammation-genetic interactions (Allen et al., 2022). Through our single cell analysis of the different tumor types, we determined that the expression of *BRAF^V600E^* or *Msh2* deletion in Min mice altered the differentiation of inflammation-induced colon tumors. We determined that the expression of mutant *BRAF* increases the differentiation of colon tumor cells and that loss of *Msh2* reduces colon tumor cell differentiation, increasing the tumor stem cell population compared to Min tumors.

The expression of mutant BRAF has been reported to increase colon epithelium differentiation (Riemer et al., 2015; Tong et al., 2017). For example, expression of *BRAF*^V600E^ has been shown to trigger intestinal stem cell differentiation (Riemer et al., 2015; Tong et al., 2017). Previously, we demonstrated that ETBF-induced *BRAF* mutant colon tumors are characterized by a mucinous phenotype (Destefano Shields et al., 2021). Here, we demonstrated that BLM tumors are more differentiated than Min tumors with an increase in enterocytes, goblet cells, and secretory-like cell populations. The increase in both goblet and secretory-like cells may contribute to the mucinous phenotype in BLM tumors. The expression of *BRAF*^V600E^ in intestinal epithelium reduces the level of intestinal stem cell markers *OLFM4* (Tong et al., 2017). We demonstrated here that the stem cells of BLM tumors exhibited low WNT pathway activity. Our data further showed that the low WNT activity in stem cells of BLM tumors is accompanied by low expression of WNT/stemness-related genes, such as *Axin2*, *Lgr5*, *Sox4*, *Id2,* and *Id1*, in comparison to Min tumors. Consistent with more differentiated BLM tumors, our trajectory analysis showed more cell directionality with a higher rate of differentiation toward differentiation lineages. Our data suggest that activation of *BRAF* shifts the stem-differentiation balance toward a differentiated cell state.

Loss of MSH2 is associated with microsatellite instability (MSI) in CRC (Armaghany et al., 2012), and we have previously shown that MSH2KO tumors had more MSI compared to Min tumors (Maiuri et al., 2017). MSI CRC is characterized by poor differentiation (H. Kim et al., 1994; Xiao et al., 2013). Consistent with this finding, our data demonstrated that MSH2KO tumors have more stem cells and less differentiated cells, such as goblet cells and enterocytes, compared to Min tumors. Additionally, MSH2KO cells had more directionality and speed toward the stem cell population. We previously showed that ETBF-induced Min and MSH2KO tumors have the same level of β-catenin (Maiuri et al., 2017). However, our scRNA-seq data revealed that MSH2KO stem cells exhibit an increase in WNT signaling pathway and WNT-related genes such as *Axin2*, *Wnt6*, and *Wnt10a* compared to Min tumor stem cells. Additionally, organoids derived from MSH2KO tumors exhibited higher expression levels of *Axin2,* a WNT target gene (Jho et al., 2002), compared to Min organoids suggesting that the difference in gene expression is intrinsic to the tumor epithelial cells and not driven by other cells in the tumor microenvironment. The WNT signaling pathway is involved in maintaining the intestinal stem cell population and increased WNT signaling activity is associated with poorly differentiated CRC (Zhao et al., 2022). Therefore, we suggest that the increased WNT signaling activity in MSH2KO contributes to fewer differentiated cells in these tumors.

We also identified known and novel regulators that might be involved in the specification of different lineages. We found that BLM tumors showed a higher level of *Cdx2*-expressing cells compared to Min tumors, which was confirmed by using tumor-derived organoids. Additionally, BLM tumor tissue sections exhibited a higher level of CDX2 protein compared to Min tumor tissue. CDX2 is an intestinal transcription factor that is involved in intestinal development (Gendron et al., 2006; Kumar et al., 2019). Loss of CDX2 in *BRAF* mutant CRC is associated with an increased stem cell population and increased oncogenic burden of the *BRAF* mutation (Sakamoto et al., 2017; Tong et al., 2017). Loss of CDX2 expression, usually through DNA methylation (J. H. Kim et al., 2013), concurrently with *BRAF* mutation is associated with poor prognosis in CRC patients (Aasebø et al., 2020; Landau et al., 2014). Interestingly, we showed that knocking down Cdx2 significantly reduced the expression of differentiated lineages-related genes such as *Muc2*, *Atoh1*, and *Guca2a* in *BRAF* mutant tumor-derived organoids. Our data suggests that CDX2 induces the differentiation of BLM tumors. Additionally, consistent with low WNT activity in BLM tumors, CDX2 has been previously shown to suppress WNT signaling activity in CRC cells (Guo et al., 2010). Interestingly, normal BLM and Min colon epithelium exhibited the same level of CDX2 protein whereas the level of CDX2 protein drastically decreased in Min colon tumors. It is possible that the expression of *BRAF*^V600E^ in Min mice maintains the expression of *Cdx2* in ETBF-induced colon tumors. Further investigation is required to study how *BRAF*^V600E^ regulates *Cdx2* expression in ETBF-induced colon tumors.

Paneth cells are known to be the source of the antimicrobial hormones, GUCA2A and REG3G, which play an essential role in intestinal homeostasis (Hoffsten et al., 2023; Shin et al., 2023). We previously demonstrated through bulk RNA-seq that ETBF induces an increase in *Reg3g* expression in BLM tumors compared to Min tumors (Destefano Shields et al., 2021). Here we determined that as expected *Guca2a* and *Reg3g* were mainly expressed by Paneth cells in Min tumors. However, they were expressed in both secretory cells and mainly in enterocytes in BLM tumors. Our findings suggest that BLM tumors have altered colon epithelial function allowing for the production and possible secretion of antimicrobial hormones in multiple differentiated lineages in response to ETBF colonization. Additionally, we found that BLM tumors had a significant increase in the secretory-like population, which was characterized by the expression of *Saa3.* SAA3 was previously reported to ameliorate dextran sodium sulfate (DSS)-induced colitis and maintain the expression of antimicrobial peptides *Reg3g* and *Reg3b* (G. Zhang et al., 2018) suggesting that secretory-like cells might be linked to the increased expression of the anti-microbial peptides in BLM tumors. It is also possible that the increased differentiation in BLM tumors is accompanied by better colon homeostasis through increasing the expression of anti-microbial peptides to maintain colon integrity in response to inflammation.

Our RNA velocity suggested *Ndrg1* as a potential additional driver for BLM tumor differentiation. NDRG1 is associated with differentiation in other cancers and cell types (Joshi et al., 2022). Additionally, NDRG1 inhibits WNT activity by preventing the nuclear localization of β-catenin (Chekmarev et al., 2021). *Ndrg1* showed high velocity expression in the stem cell population of BLM tumors, while it was absent in Min tumor stem cells. This result suggests that low WNT activity in BLM tumors may be due to the high velocity level of *Ndrg1* in these tumors. Other regulators for colon tumor differentiation were identified such as *Styl2*, *Kcnb2*, *Nrg1*, and *Mpp5*. Future work will be required to extend our findings by exploring how Ndrg1 mechanistically regulates differentiation in BLM colon tumors.

Overall, we propose a model where increased expression of WNT/stemness-related genes and increased activity of WNT signaling in MSH2KO tumors maintain the stem cell population resulting in more cycling stem cells in MSH2KO tumors and reducing their tendency to differentiate toward different lineages. However, low WNT activity in BLM tumors increases the differentiation potential of BLM colon tumors. Furthermore, the increased expression of *Cdx2*, and other differentiation-driving transcription factors like *Klf4* and the WNT antagonist, *Ndrg1*, in BLM tumors push the stem cells toward the various differentiated lineages (Supplemental Fig. S6E). Determining how the differentiation of inflammation-associated colon tumor epithelial cells is regulated in different genetic backgrounds will lead to a greater understanding of tumor epithelial cell biology that has the potential to alter therapy response.

## Methods

### Animal model

Min^ApcΔ716/+^ mice were handled and inoculated with Enterotoxigenic Bacteroides *fragilis* (ETBF) as in Wu et al (Wu et al., 2009). Msh2^l/l^VC are a result of crossing B6.Cg-MSH2^tm2.1Rak^/J (The Jackson Laboratory; RRID:IMSR_JAX:016231) and B6.Cg-Tg(Vil1-cre) 997Gum/J mice (RRID:IMSR_JAX:004586) to create mice homozygous for MSH2^tm2.1Rak^ and expressing the Vil1-cre transgene. Msh2^l/l^VC/Min (MSH2KO) mice are the result of crossing Msh2^l/l^VC and Min^ApcΔ716/+^ mice. Mice containing Loxp flanked *BRAF*^F-V600E^ (B6.129P2(Cg)-*Braf^tm1Mmcm/J^*; RRID: IMSR_JAX:017837) and leucine-rich repeat-containing G protein-coupled receptor 5 (*Lgr5*) CreERT2 knock-in (*Lgr5*^tm1(Cre/ERT2)Cle^; RRID: IMSR_JAX:008875) were crossed with Min^ApcΔ716/+^ mice to produce *BRAF*^F-V600E^*Lgr5*^tm1(Cre/ERT2)Cle^ Min^ApcΔ716/+^ (BLM) mice. Recombination in mice bearing Lgr5^Cre^ was induced with tamoxifen, as in Barker et al (Barker et al., 2007), at 4 weeks of age. All mice were bred and maintained in a specific pathogen-free barrier facility, both males and females were used for all experiments, and mice of different genotypes were cohoused. 7-8 weeks post-ETBF colonization mice were euthanized, and individual tumors were removed from dissected colons with the aid of a dissecting microscope, pooled, digested and used for scRNA-seq and organoid derivation as indicated below. Alternatively, dissected colons were swiss rolled, fixed in 10% formalin for 48H and paraffin embedded (FFPE). All mouse experiments were covered under an approved Indiana University Bloomington Animal Care and Use Committee protocol, in accordance with the Association for Assessment and Accreditation of Laboratory Animal Care International.

### Tumor digestion and single cell RNA sequencing

To obtain single cell solutions of tumor cells, pooled distal (Min, MSH2KO), distal and proximal (BLM) or proximal (BLM2) tumors were washed in HBSS, incubated in 0.25% trypsin-EDTA for 10 minutes at 37°C, followed by the addition of FBS to inactivate the trypsin. Next, the tumors were digested by incubation in liberase TM (0.05 mg/ml; Roche, #05401119001)+DNAse (0.2 mg/ml) in DMEM at 37°C for 2H while rotating. Following washing in HBSS and an additional incubation in 0.25% trypsin-EDTA for 10 minutes at 37°C, cells were resuspended in DMEM+10%FBS and filtered through a 40 μM mesh cell strainer. Cells were then washed with DPBS+0.1% BSA and viable cells were counted. Cells were resuspended at 1,000 cell per μl in DPBS+0.1% BSA. All single cell preparations used for sequencing had a viability of > 80%. 10,000 cells per sample were targeted for input to the 10X Genomics Chromium system using the Chromium Next GEM Single Cell 30 Kit v3.1 at the Indiana University School of Medicine (IUSM) Center for Medical Genomics core. The libraries were sequenced at the IUSM Center for Medical Genomics using a NovaSeq 6000 with a NovaSeq S2 reagent kit v1.0 (100 cycles) with approximately 450 million read pairs per sample. The remaining cells from each single cell preparation were used to derive organoids.

### Cells and Organoids

HEK293T cell and organoids were maintained in a humidified atmosphere at 37 C with 5% CO2. HEK293Tcells were cultured in DMEM 1X (Corning, #10-013-CV) with 10% FBS (Corning, #35-015-CV) without antibiotics. Organoids were derived from disassociated colon tumors (BLM, BLM2, Min, and MSH2KO) and grown in growth factor reduced Matrigel plus organoid media (advanced DMEM/F12 (Gibco, #12634-010) supplemented with EGF (R&D Systems, #236-EG), Noggin (R&D systems, #6057-NG), N2 supplement (17502048, Fisher), B27 supplement (Fisher, #17504044), HEPES, and Penn/Strep) as in (Maiuri et al., 2018).

### Generation of stable knockdown organoids

For knockdown of Cdx2 (Sigma, NM_007673, #TRCN0000055393), Ndrg1 (Sigma, NM_008681, #TRCN0000238073), and empty vector (EV) TRC2 (Sigma, #SHC201), the lentiviral shRNA knockdown protocol from The RNAi Consortium Broad Institute was used as in (Ghobashi et al., 2023). Briefly, 4 x 10^5^ HEK293T cells were plated on day 1 in DMEM 1X containing 10% FBS. On day 2, cells were transfected with shRNA of interest, EV control, and packaging plasmids. On day 3, the media was replaced with fresh DMEM containing 10% FBS. Approximately 24 hours later, media containing lentiviral particles was collected, and fresh DMEM + 10% FBS was added. The added media were collected 24 hours later and pooled with media harvested on day 4. The pooled media was then filtered using a 0.45 mm filter and concentrated using a Lenti-X™ Concentrator (Takara, #631232). To perform the knockdown, concentrated virus plus polybrene was added to the organoids. Cells were treated with puromycin (1 mg/mL) (Sigma-Aldrich, #P8833) after 24 hours to select for knockdown organoids.

### RNA isolation and Gene expression

RNA was prepared from organoids using TRIzol followed by cleanup with RNAeasy micro kit (Qiagen, #74004) as per the manufacturer’s protocol. Maxima first strand cDNA synthesis kit (Thermo Fisher, #K1642) for quantitative reverse transcription PCR was used to synthesize cDNA. qPCR was done using TaqMan assays (see Supplementary Table S3 for assays used). Expression of candidate genes was normalized to the expression of a housekeeping gene (*Ppia*).

### Immunohistochemistry (IHC)

CDX2 and SAA3 were detected by IHC on 8-μm FFPE colon tissue samples following unmasking in TRIS/EDTA buffer (CDX2) or citrate buffer (SAA3). Anti-CDX2 (ab76541) and anti-SAA3 antibody (ab231680) were applied at a dilution of 1:50, followed by rabbit HRP SignalStain Boost (Cell Signaling Technology, #8125), rat HRP SignalStain Boost (Cell Signaling Technology, #72838), respectively, and DAB substrate (CST, #8059). Slides were counterstained in hematoxylin. Tumors stained for CDX2 or SAA3 were scored from 0 to 4. 0: no staining, 1: >=10% of the tumor epithelial cells were positively stained, 2: 11-33% stained, 3: 34-50% stained, and 4: >50% stained.

### Immunofluorescence and imaging

FFPE colon tissue samples were unmasked in citrate buffer, blocked, and then incubated with anti-SAA3 (ab231680, 1:50), and anti-E-cadherin (cell signaling technology, #3195, 1:200) in 1% BSA in PBST overnight at 4°C. Then, the tissues were incubated with Alexa Fluor (AF) -conjugated secondary antibodies (anti-rat AF488, CST, #4416; anti-rabbit AF594, CST, #8889) for 2H at room temperature. Images were taken using an EVOS FL Auto microscope (Life Technologies)

### Statistical analysis

Expression data and IHC are presented as the mean +/- SEM. These data are evaluated by a one-tailed t-test and considered statistically significant with a P < 0.05.

### Computational analysis

#### Single-cell data pre-processing and QC

Read alignment and gene-expression quantification of mouse scRNA-seq data was performed using the CellRanger Count pipeline (version 3.1.0, 10X Genomics). The CellRanger pre-built mouse reference package was used for read alignment (mm10). The filtered feature matrices output was then used to create Seurat object using Seurat package v4.3.0.1 (Hao et al., 2021). Cells were filtered to include only cells with no more than 20% mitochondrial gene expression and doublets were removed using DoubletFinder v2.0 (McGinnis et al., 2019). The data were normalized, and highly variable genes were identified and scaled using *SCTransform*. Next, dimensionality reduction by principal components (PCs) was calculated using *RunPCA* and to estimate the significant number of PCs to be used *ElbowPlot* function was used. Next, the uniform manifold approximation and projection (UMAP) embedding were calculated and visualized using *RunUMAP* and *DimPlot*. Unsupervised Louvain clustering was carried out using *FindNeighbors* and *FindClusters*. Differentially expressed genes were then defined with *FindallMarkers* with Wilcox test.

### Data integration with batch correction

In our analysis, we used Seurat v4.3.0.1 to perform batch-effect correction. 3000 highly variable genes were defined within the 4 mouse samples with the Seurat *FindVariableFeatures* function. We also identified unsupervised integration “anchors” for similar cell states using shared nearest neighbor graphs (*FindIntegrationAnchors*), and then integrated our 4 different datasets using these anchors using *IntegrateData*. The output was then transformed into principal component analysis (PCA) space for further evaluation and visualization.

### Subsetting and visualizing epithelial data

To obtain a Seurat object containing only the epithelial cell type of the integrated data, the ‘‘subset’’ function was used. The subset Seurat object goes through Seurat filtering, normalization, and integration workflows as described above. Proportional difference in epithelial cell populations between two samples was computed using the R package scProportion (https://github.com/rpolicastro/scProportionTest/releases/tag/v1.0.0)(Miller et al., 2021). Gene set enrichment scores for single cells were computed using escape (v1.12.0)(Borcherding et al., 2021). Diffusion map, diffusion pseudotime, and cell density were computed using *sc.tl.diffmap*, *sc.tl.dpt*, and *sc.pl.embedding_density* respectively, which are implemented through Scanpy (v1.9.6)(Wolf et al., 2018). The stem cell was annotated manually as a root cell before computing diffusion pseudotime.

### Gene Ontology enrichment analysis

Gene Ontology (GO) enrichment was performed using Metascape (https://metascape.org/gp/index.html)(Zhou et al., 2019).

### RNA velocity

Spliced/unspliced expression matrices were generated as loom files using Velocyto (La Manno et al., 2018). Seurat objects were converted into AnnData objects containing the corrected counts, clusters, and UMAP embeddings. Then, the loom files were merged with the AnnData objects and loaded into scVelo (v0.2.1)(Bergen et al., 2020), the ratio of spliced to unspliced reads per cluster was found, and cell velocities were computed. All functions were run with default settings unless otherwise stated. The *scvelo.pp.filter_and_normalize* argument ‘n_top_genes’ was set to 3000, and the ‘n_npcs’ and ‘n_neighbors’ arguments of *scvelo.pp.momentum* were both set to 30. The velocity cell arrows were made with the *scvelo.pl.velocity_embedding* function. The top velocity genes per cluster were discovered using *scvelo.tl.rank_velocity_genes*, and plotted using *scvelo.pl.velocity*. *scvelo.tl.velocity_confidence* generated the velocity confidence and length values, and the results were plotted using *scvelo.pl.scatter*. Cell cycle signatures were computed using *scv.tl.score_genes_cell_cycle* and plotted using *scvelo.pl.scatter*. The RNA-velocity analysis was extended through calculating RNA splicing kinetics using dynamic model using *scv.tl.recover_dynamics* and scv.tl.*velocity*(mode=’dynamical’). cluster-specific identification of potential drivers was discovered using *scv.tl.rank_dynamical_genes*. By applying *scv.tl.differential_kinetic_test* to the dynamic model, we were able to discover which cluster exhibited significant RNA splicing kinetics for *Guca2a* transcript, and the results were plotted using *scvelo.pl.velocity*.

### Simulated gene perturbation

Simulated gene perturbation was performed using CellOracle (Kamimoto et al., 2023). We used gene-regulatory networks (GRN) from mouse scATAC-seq data using *co.data.load_mouse_scATAC_atlas_base_GRN*. Then GRN data and BLM2 gene expression matrix were loaded into the CellOracle object using *co.import_TF_data* and *co.import_anndata_as_raw_count*, respectively. We constructed a cluster-specific GRN for all clusters using *oc.get_links* and kept only network edges with p-value <=0.01. To simulate gene overexpression or knockout, we perturb the gene expression to 1 or 0, respectively in the *oc.simulate_shift* function.

## Acknowledgments

We would like to thank the Indiana University School of Medicine (IUSM) Center for Medical Genomics core for their assistance with performing the scRNA-seq and the Indiana Center for Musculoskeletal Health Histology Core for their assistance with embedding and sectioning tissue samples. This work was in part supported by a Research Enhancement Grant [to H. M. O’Hagan] from the Indiana University School of Medicine (IUSM) and pilot funding [to H. M. O’Hagan] from the IU Simon Comprehensive Cancer Center (IUSCCC) Tumor Microenvironment & Metastasis Program and the IUSCCC P30 Support Grant (P30CA082709). The content is solely the responsibility of the authors and does not necessarily represent the official views of the NIH or IUSM. Additional pilot funding was provided by the Van Andel Institute through the Van Andel Institute - Stand Up to Cancer Epigenetics Dream Team and the NCI SPORE Project, Epigenetic Therapies – New Approaches Developmental Research Program [to H. M. O’Hagan]. Stand Up to Cancer is a division of the Entertainment Industry Foundation, administered by AACR. A. Ghobashi and C. Ladaika were supported by the Doane and Eunice Dahl Wright Fellowship generously provided by Ms. Imogen Dahl.

## Author Contributions

A.G. Conceptualization, formal analysis, computational analysis, validation, investigation, visualization, methodology, writing–original draft, writing–review and editing. R.L. Investigation. C.L. Investigation. H.O. Conceptualization, resources, supervised the study, contributed to the conception and design, and helped write and revise the manuscript. All authors reviewed and approved the final manuscript.

## References

Aasebø, K., Dragomir, A., Sundström, M., Mezheyeuski, A., Edqvist, P. H., Eide, G. E., Ponten, F., Pfeiffer, P., Glimelius, B., & Sorbye, H. (2020). CDX2: A Prognostic Marker in Metastatic Colorectal Cancer Defining a Better BRAF Mutated and a Worse KRAS Mutated Subgroup. Frontiers in Oncology, 10(February). 10.3389/fonc.2020.00008

Allen, J., Rosendahl Huber, A., Pleguezuelos-Manzano, C., Puschhof, J., Wu, S., Wu, X., Boot, C., Saftien, A., O’Hagan, H. M., Wang, H., van Boxtel, R., Clevers, H., & Sears, C. L. (2022). Colon Tumors in Enterotoxigenic Bacteroides fragilis (ETBF)-Colonized Mice Do Not Display a Unique Mutational Signature but Instead Possess Host-Dependent Alterations in the APC Gene. Microbiology Spectrum, 10(3). 10.1128/spectrum.01055-22

Armaghany, T., Wilson, J. D., Chu, Q., & Mills, G. (2012). Genetic alterations in colorectal cancer. Gastrointestinal Cancer Research: GCR, 5(1), 19–27.

Barker, N., van Es, J. H., Kuipers, J., Kujala, P., van den Born, M., Cozijnsen, M., Haegebarth, A., Korving, J., Begthel, H., Peters, P. J., & Clevers, H. (2007). Identification of stem cells in small intestine and colon by marker gene Lgr5. Nature, 449(7165), 1003–1007. 10.1038/nature06196

Bergen, V., Lange, M., Peidli, S., Wolf, F. A., & Theis, F. J. (2020). Generalizing RNA velocity to transient cell states through dynamical modeling. Nature Biotechnology, 38(12), 1408–1414. 10.1038/s41587-020-0591-3

Boleij, A., Hechenbleikner, E. M., Goodwin, A. C., Badani, R., Stein, E. M., Lazarev, M. G., Ellis, B., Carroll, K. C., Albesiano, E., Wick, E. C., Platz, E. A., Pardoll, D. M., & Sears, C. L. (2015). The bacteroides fragilis toxin gene is prevalent in the colon mucosa of colorectal cancer patients. Clinical Infectious Diseases, 60(2), 208–215. 10.1093/cid/ciu787

Borcherding, N., Vishwakarma, A., Voigt, A. P., Bellizzi, A., Kaplan, J., Nepple, K., Salem, A. K., Jenkins, R. W., Zakharia, Y., & Zhang, W. (2021). Mapping the immune environment in clear cell renal carcinoma by single-cell genomics. Communications Biology, 4(1), 1–11. 10.1038/s42003-020-01625-6

Cai, K., El-Merahbi, R., Loeffler, M., Mayer, A. E., & Sumara, G. (2017). Ndrg1 promotes adipocyte differentiation and sustains their function. Scientific Reports, 7(1), 1–9. 10.1038/s41598-017-07497-x

Chang, S. K., Dohrman, A. F., Basbaum, C. B., Ho, S. B., Tsuda, T., Toribara, N. W., Gum, J. R., & Kim, Y. S. (1994). Localization of mucin (MUC2 and MUC3) messenger RNA and peptide expression in human normal intestine and colon cancer. Gastroenterology, 107(1), 28–36. 10.1016/0016-5085(94)90057-4

Chekmarev, J., Azad, M. G., & Richardson, D. R. (2021). The Oncogenic Signaling Disruptor, NDRG1: Molecular and Cellular Mechanisms of Activity. Cells, 10(9). 10.3390/cells10092382

Chiba, T., Marusawa, H., & Ushijima, T. (2012). Inflammation-associated cancer development in digestive organs: Mechanisms and roles for genetic and epigenetic modulation. Gastroenterology, 143(3), 550–563. 10.1053/j.gastro.2012.07.009

Clevers, H. (2013). The intestinal crypt, a prototype stem cell compartment. Cell, 154(2), 274. 10.1016/j.cell.2013.07.004

Dagogo-Jack, I., & Shaw, A. T. (2018). Tumour heterogeneity and resistance to cancer therapies. Nature Reviews Clinical Oncology, 15(2), 81–94. 10.1038/nrclinonc.2017.166

Destefano Shields, C. E., White, J. R., Chung, L., Wenzel, A., Hicks, J. L., Tam, A. J., Chan, J. L., Dejea, C. M., Fan, H., Michel, J., Maiuri, A. R., Sriramkumar, S., Podicheti, R., Rusch, D. B., Wang, H., De Marzo, A. M., Besharati, S., Anders, R. A., Baylin, S. B., … Sears, C. L. (2021). Bacterial-driven inflammation and mutant braf expression combine to promote murine colon tumorigenesis that is sensitive to immune checkpoint therapy. Cancer Discovery, 11(7), 1792–1807. 10.1158/2159-8290.CD-20-0770

Fan, X., Jin, Y., Chen, G., Ma, X., & Zhang, L. (2021). Gut Microbiota Dysbiosis Drives the Development of Colorectal Cancer. Digestion, 102(4), 508–515. 10.1159/000508328

Gendron, F. P., Mongrain, S., Laprise, P., McMahon, S., Dubois, C. M., Blais, M., Asselin, C., & Rivard, N. (2006). The CDX2 transcription factor regulates furin expression during intestinal epithelial cell differentiation. American Journal of Physiology - Gastrointestinal and Liver Physiology, 290(2), 310–318. 10.1152/ajpgi.00217.2005

Ghobashi, A. H., Vuong, T. T., Kimani, J. W., Ladaika, C. A., Hollenhorst, P. C., & O’Hagan, H. M. (2023). Activation of AKT induces EZH2-mediated β-catenin trimethylation in colorectal cancer. IScience, 26(9), 107630. 10.1016/j.isci.2023.107630

Gorman, H., Moreau, F., Dufour, A., & Chadee, K. (2023). IgGFc-binding protein and MUC2 mucin produced by colonic goblet-like cells spatially interact non-covalently and regulate wound healing. Frontiers in Immunology, 14(June), 1–25. 10.3389/fimmu.2023.1211336

Guo, R. J., Funakoshi, S., Lee, H. H., Kong, J., & Lynch, J. P. (2010). The intestine-specific transcription factor Cdx2 inhibits β-catenin/TCF transcriptional activity by disrupting the β-catenin-TCF protein complex. Carcinogenesis, 31(2), 159–166. 10.1093/carcin/bgp213

Haber, A. L., Biton, M., Rogel, N., Herbst, R. H., Shekhar, K., Smillie, C., Burgin, G., Delorey, T. M., Howitt, M. R., Katz, Y., Tirosh, I., Beyaz, S., Dionne, D., Zhang, M., Raychowdhury, R., Garrett, W. S., Rozenblatt-Rosen, O., Shi, H. N., Yilmaz, O., … Regev, A. (2017). A single-cell survey of the small intestinal epithelium. Nature, 551(7680), 333–339. 10.1038/nature24489

Haghverdi, L., Büttner, M., Wolf, F. A., Buettner, F., & Theis, F. J. (2016). Diffusion pseudotime robustly reconstructs lineage branching. Nature Methods, 13(10), 845–848. 10.1038/nmeth.3971

Hao, Y., Hao, S., Andersen-Nissen, E., Mauck, W. M., Zheng, S., Butler, A., Lee, M. J., Wilk, A. J., Darby, C., Zager, M., Hoffman, P., Stoeckius, M., Papalexi, E., Mimitou, E. P., Jain, J., Srivastava, A., Stuart, T., Fleming, L. M., Yeung, B., … Satija, R. (2021). Integrated analysis of multimodal single-cell data. Cell, 184(13), 3573–3587.e29. 10.1016/j.cell.2021.04.048

Hickey, J. W., Becker, W. R., Nevins, S. A., Horning, A., Perez, A. E., Zhu, C., Zhu, B., Wei, B., Chiu, R., Chen, D. C., Cotter, D. L., Esplin, E. D., Weimer, A. K., Caraccio, C., Venkataraaman, V., Schürch, C. M., Black, S., Brbić, M., Cao, K., … Snyder, M. (2023). Organization of the human intestine at single-cell resolution. Nature, 619(7970), 572–584. 10.1038/s41586-023-05915-x

Hoffsten, A., Lilja, H. E., Mobini-Far, H., Sindelar, R., & Markasz, L. (2023). Paneth cell proteins DEFA6 and GUCA2A as tissue markers in necrotizing enterocolitis. European Journal of Pediatrics, 182(6), 2775–2784. 10.1007/s00431-023-04907-3

Jho, E., Zhang, T., Domon, C., Joo, C.-K., Freund, J.-N., & Costantini, F. (2002). Wnt/β-Catenin/Tcf Signaling Induces the Transcription of Axin2, a Negative Regulator of the Signaling Pathway. Molecular and Cellular Biology, 22(4), 1172–1183. 10.1128/mcb.22.4.1172-1183.2002

Joshi, V., Lakhani, S. R., & McCart Reed, A. E. (2022). NDRG1 in Cancer: A Suppressor, Promoter, or Both? Cancers, 14(23). 10.3390/cancers14235739

Kamimoto, K., Stringa, B., Hoffmann, C. M., Jindal, K., Solnica-Krezel, L., & Morris, S. A. (2023). Dissecting cell identity via network inference and in silico gene perturbation. Nature, 614(7949), 742–751. 10.1038/s41586-022-05688-9

Kim, H., Jen, J., Vogelstein, B., & Hamilton, S. R. (1994). Clinical and pathological characteristics of sporadic colorectal carcinomas with DNA replication errors in microsatellite sequences. The American Journal of Pathology, 145(1), 148–156.

Kim, J. H., Rhee, Y.-Y., Bae, J. M., Cho, N.-Y., & Kang, G. H. (2013). Loss of CDX2/CK20 expression is associated with poorly differentiated carcinoma, the CpG island methylator phenotype, and adverse prognosis in microsatellite-unstable colorectal cancer. The American Journal of Surgical Pathology, 37(10), 1532–1541. 10.1097/PAS.0b013e31829ab1c1

Kumar, N., Tsai, Y.-H., Chen, L., Zhou, A., Banerjee, K. K., Saxena, M., Huang, S., Toke, N. H., Xing, J., Shivdasani, R. A., Spence, J. R., & Verzi, M. P. (2019). The lineage-specific transcription factor CDX2 navigates dynamic chromatin to control distinct stages of intestine development. Development (Cambridge, England), 146(5). 10.1242/dev.172189

La Manno, G., Soldatov, R., Zeisel, A., Braun, E., Hochgerner, H., Petukhov, V., Lidschreiber, K., Kastriti, M. E., Lönnerberg, P., Furlan, A., Fan, J., Borm, L. E., Liu, Z., van Bruggen, D., Guo, J., He, X., Barker, R., Sundström, E., Castelo-Branco, G., … Kharchenko, P. V. (2018). RNA velocity of single cells. Nature, 560(7719), 494–498. 10.1038/s41586-018-0414-6

Landau, M. S., Kuan, S. F., Chiosea, S., & Pai, R. K. (2014). BRAF-mutated microsatellite stable colorectal carcinoma: An aggressive adenocarcinoma with reduced CDX2 and increased cytokeratin 7 immunohistochemical expression. Human Pathology, 45(8), 1704–1712. 10.1016/j.humpath.2014.04.008

Li, X., Liu, G., & Wu, W. (2021). Recent advances in Lynch syndrome. Experimental Hematology and Oncology, 10(1), 4–11. 10.1186/s40164-021-00231-4

Liang, X., Xie, J., Liu, H., Zhao, R., Zhang, W., Wang, H., Pan, H., Zhou, Y., & Han, W. (2022). STIM1 Deficiency In Intestinal Epithelium Attenuates Colonic Inflammation and Tumorigenesis by Reducing ER Stress of Goblet Cells. Cmgh, 14(1), 193–217. 10.1016/j.jcmgh.2022.03.007

Lo, Y.-H., Chung, E., Li, Z., Wan, Y.-W., Mahe, M. M., Chen, M.-S., Noah, T. K., Bell, K. N., Yalamanchili, H. K., Klisch, T. J., Liu, Z., Park, J.-S., & Shroyer, N. F. (2017). Transcriptional Regulation by ATOH1 and its Target SPDEF in the Intestine. Cellular and Molecular Gastroenterology and Hepatology, 3(1), 51–71. 10.1016/j.jcmgh.2016.10.001

Lucafò, M., Curci, D., Franzin, M., Decorti, G., & Stocco, G. (2021). Inflammatory Bowel Disease and Risk of Colorectal Cancer: An Overview From Pathophysiology to Pharmacological Prevention. Frontiers in Pharmacology, 12(October), 1–9. 10.3389/fphar.2021.772101

Maiuri, A. R., Li, H., Stein, B. D., Tennessen, J. M., & O’Hagan, H. M. (2018). Inflammation-induced DNA methylation of DNA polymerase gamma alters the metabolic profile of colon tumors. Cancer & Metabolism, 6(1), 1–11. 10.1186/s40170-018-0182-7

Maiuri, A. R., Peng, M., Sriramkumar, S., Kamplain, C. M., DeStefano Shields, C. E., Sears, C. L., & O’Hagan, H. M. (2017). Mismatch repair proteins initiate epigenetic alterations during inflammation-driven tumorigenesis. Cancer Research, 77(13), 3467–3478. 10.1158/0008-5472.CAN-17-0056

McGinnis, C. S., Murrow, L. M., & Gartner, Z. J. (2019). DoubletFinder: Doublet Detection in Single-Cell RNA Sequencing Data Using Artificial Nearest Neighbors. Cell Systems, 8(4), 329–337.e4. 10.1016/j.cels.2019.03.003

Miller, S. A., Policastro, R. A., Sriramkumar, S., Lai, T., Huntington, T. D., Ladaika, C. A., Kim, D., Hao, C., Zentner, G. E., & O’Hagan, H. M. (2021). LSD1 and aberrant DNA methylation mediate persistence of enteroendocrine progenitors that support BRAF-mutant colorectal cancer. Cancer Research, 81(14), 3791–3805. 10.1158/0008-5472.CAN-20-3562

Norreys, P. A., Sakabe, S., Tanaka, K. A., Youssef, A., Zepf, M., & Yamanaka, T. (1994). Tumorigenesis RAF/RAS oncogenes and mismatch-repair status. Conf. Laser Elec. Opt, 1(2), 402–403. www.nature.com/nature

Reigstad, C. S., & Bäckhed, F. (2010). Microbial regulation of SAA3 expression in mouse colon and adipose tissue. Gut Microbes, 1(1), 55–57. 10.4161/gmic.1.1.10514

Riemer, P., Sreekumar, A., Reinke, S., Rad, R., Schäfer, R., Sers, C., Bläker, H., Herrmann, B. G., & Morkel, M. (2015). Transgenic expression of oncogenic BRAF induces loss of stem cells in the mouse intestine, which is antagonized by β-catenin activity. Oncogene, 34(24), 3164–3175. 10.1038/onc.2014.247

Sakamoto, N., Feng, Y., Stolfi, C., Kurosu, Y., Green, M., Lin, J., Green, M. E., Sentani, K., Yasui, W., McMahon, M., Hardiman, K. M., Spence, J. R., Horita, N., Greenson, J. K., Kuick, R., Cho, K. R., & Fearon, E. R. (2017). BRAFV600E cooperates with CDX2 inactivation to promote serrated colorectal tumorigenesis. ELife, 6, 1–25. 10.7554/eLife.20331

Schatoff, E. M., Leach, B. I., & Dow, L. E. (2017). Wnt Signaling and Colorectal Cancer. Current Colorectal Cancer Reports, 13(2), 101–110. 10.1007/s11888-017-0354-9

Shin, J. H., Bozadjieva-Kramer, N., & Seeley, R. J. (2023). Reg3γ: current understanding and future therapeutic opportunities in metabolic disease. Experimental & Molecular Medicine, 55(8), 1672– 1677. 10.1038/s12276-023-01054-5

Siegel, R. L., Miller, K. D., Wagle, N. S., & Jemal, A. (2023). Cancer statistics, 2023. CA: A Cancer Journal for Clinicians, 73(1), 17–48. 10.3322/caac.21763

Tabernero, J., Ros, J., & Élez, E. (2022). The Evolving Treatment Landscape in BRAF-V600E –Mutated Metastatic Colorectal Cancer. American Society of Clinical Oncology Educational Book, 42, 254–263. 10.1200/edbk_349561

Tong, K., Pellón-Cárdenas, O., Sirihorachai, V. R., Warder, B. N., Kothari, O. A., Perekatt, A. O., Fokas, E. E., Fullem, R. L., Zhou, A., Thackray, J. K., Tran, H., Zhang, L., Xing, J., & Verzi, M. P. (2017). Degree of Tissue Differentiation Dictates Susceptibility to BRAF-Driven Colorectal Cancer. Cell Reports, 21(13), 3833–3845. 10.1016/j.celrep.2017.11.104

Ulger Toprak, N., Yagci, A., Gulluoglu, B. M., Akin, M. L., Demirkalem, P., Celenk, T., & Soyletir, G. (2006). A possible role of Bacteroides fragilis enterotoxin in the aetiology of colorectal cancer. Clinical Microbiology and Infection, 12(8), 782–786. 10.1111/j.1469-0691.2006.01494.x

Viljoen, K. S., Dakshinamurthy, A., Goldberg, P., & Blackburn, J. M. (2015). Quantitative profiling of colorectal cancer-associated bacteria reveals associations between Fusobacterium spp., enterotoxigenic Bacteroides fragilis (ETBF) and clinicopathological features of colorectal cancer. PLoS ONE, 10(3), 1–21. 10.1371/journal.pone.0119462

Waarts, M. R., Stonestrom, A. J., Park, Y. C., & Levine, R. L. (2022). Targeting mutations in cancer. The Journal of Clinical Investigation, 132(8). 10.1172/JCI154943

Watari, K., Shibata, T., Nabeshima, H., Shinoda, A., Fukunaga, Y., Kawahara, A., Karasuyama, K., Fukushi, J. I., Iwamoto, Y., Kuwano, M., & Ono, M. (2016). Impaired differentiation of macrophage lineage cells attenuates bone remodeling and inflammatory angiogenesis in Ndrg1 deficient mice. Scientific Reports, 6(September 2015), 1–14. 10.1038/srep19470

Wolf, F. A., Angerer, P., & Theis, F. J. (2018). SCANPY: large-scale single-cell gene expression data analysis. Genome Biology, 19(1), 15. 10.1186/s13059-017-1382-0

Wu, S., Rhee, K.-J., Albesiano, E., Rabizadeh, S., Wu, X., Yen, H.-R., Huso, D. L., Brancati, F. L., Wick, E., McAllister, F., Housseau, F., Pardoll, D. M., & Sears, C. L. (2009). A human colonic commensal promotes colon tumorigenesis via activation of T helper type 17 T cell responses. Nature Medicine, 15(9), 1016–1022. 10.1038/nm.2015

Xiao, H., Yoon, Y. S., Hong, S.-M., Roh, S. A., Cho, D.-H., Yu, C. S., & Kim, J. C. (2013). Poorly differentiated colorectal cancers: correlation of microsatellite instability with clinicopathologic features and survival. American Journal of Clinical Pathology, 140(3), 341–347. 10.1309/AJCP8P2DYNKGRBVI

Yu, T., Chen, X., Zhang, W., Li, J., Xu, R., Wang, T. C., Ai, W., & Liu, C. (2012). Krüppel-like factor 4 regulates intestinal epithelial cell morphology and polarity. PLoS ONE, 7(2), 1–9. 10.1371/journal.pone.0032492

Zhang, G., Liu, J., Wu, L., Fan, Y., Sun, L., Qian, F., Chen, D., & Ye, R. D. (2018). Elevated expression of serum amyloid A 3 protects colon epithelium against acute injury through TLR2-dependent induction of neutrophil IL-22 expression in a mouse model of colitis. Frontiers in Immunology, 9(JUN), 1–11. 10.3389/fimmu.2018.01503

Zhang, L., & Shay, J. W. (2017). Multiple Roles of APC and its Therapeutic Implications in Colorectal Cancer. Journal of the National Cancer Institute, 109(8), 1–10. 10.1093/jnci/djw332

Zhao, H., Ming, T., Tang, S., Ren, S., Yang, H., Liu, M., Tao, Q., & Xu, H. (2022). Wnt signaling in colorectal cancer: pathogenic role and therapeutic target. Molecular Cancer, 21(1), 1–34. 10.1186/s12943-022-01616-7

Zhou, Y., Zhou, B., Pache, L., Chang, M., Khodabakhshi, A. H., Tanaseichuk, O., Benner, C., & Chanda, S. K. (2019). Metascape provides a biologist-oriented resource for the analysis of systems-level datasets. Nature Communications, 10(1). 10.1038/s41467-019-09234-6

