## Supplemental materials for "Single-cell profiling reveals the impact of genetic alterations on the differentiation of inflammation-induced colon tumors"

**Inventory of Supplementary Materials**

**Supplementary Figures**

Supplemental Figure S1. Violinplot of normalized gene expression of epithelial, fibroblast, lymphoid, and myeloid cell gene markers in the combined BLM, BLM2, Min, and MSH2KO scRNA-seq data.

Supplemental Figure S2. Heatmap of normalized gene expression of gene markers for stem cells, enterocyte cells (EC), transit-amplifying cells (TA), EC/brush border, goblet cells (GC), and Paneth cells (PC), and secretory-like cells in combined BLM, BLM2, Min, and MSH2KO colon epithelial tumors. (**Supplemental_Fig S2.pdf**)

Supplemental Figure S3. Featureplot of normalized gene expression of different genes in BLM, BLM2, Min, and MSH2KO colon epithelial tumors.

Supplemental Figure S5. Stem cells of BLM and MSH2KO colon epithelial tumors have low and high WNT signaling activity, respectively.

Supplemental Figure S6. *Cdx2* and *Ndrg1* are involved in BLM colon tumor epithelial cell differentiation.

**Supplementary Tables**

Table S1. Supplemental_Table_S1.xls.

Table S2. Supplemental_Table_S2.xls

Table S3. Taqman qPCR assays

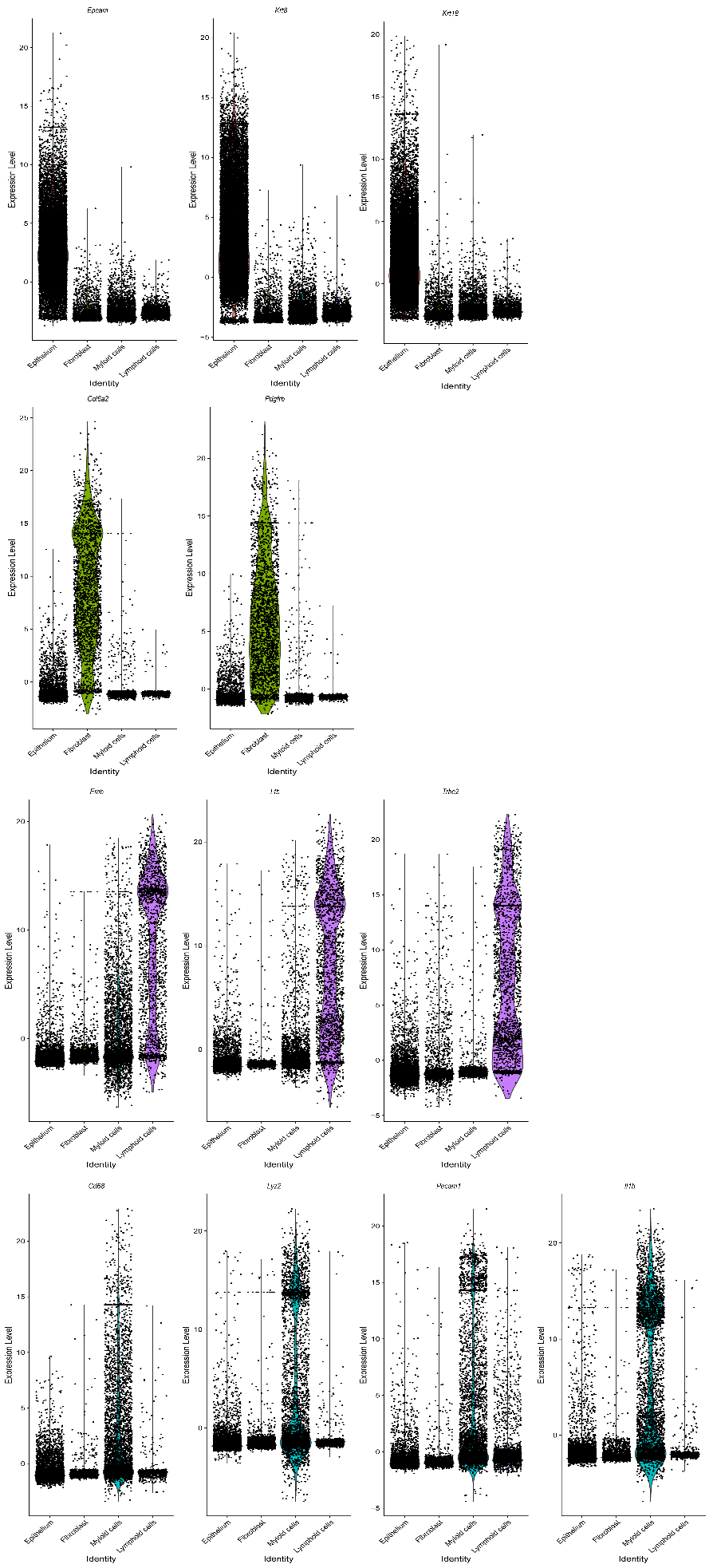

Figure S1. Violinplot of normalized gene expression of epithelial, fibroblast, lymphoid, and myeloid cell gene markers in the combined BLM, BLM2, Min, and MSH2KO scRNA-seq data.

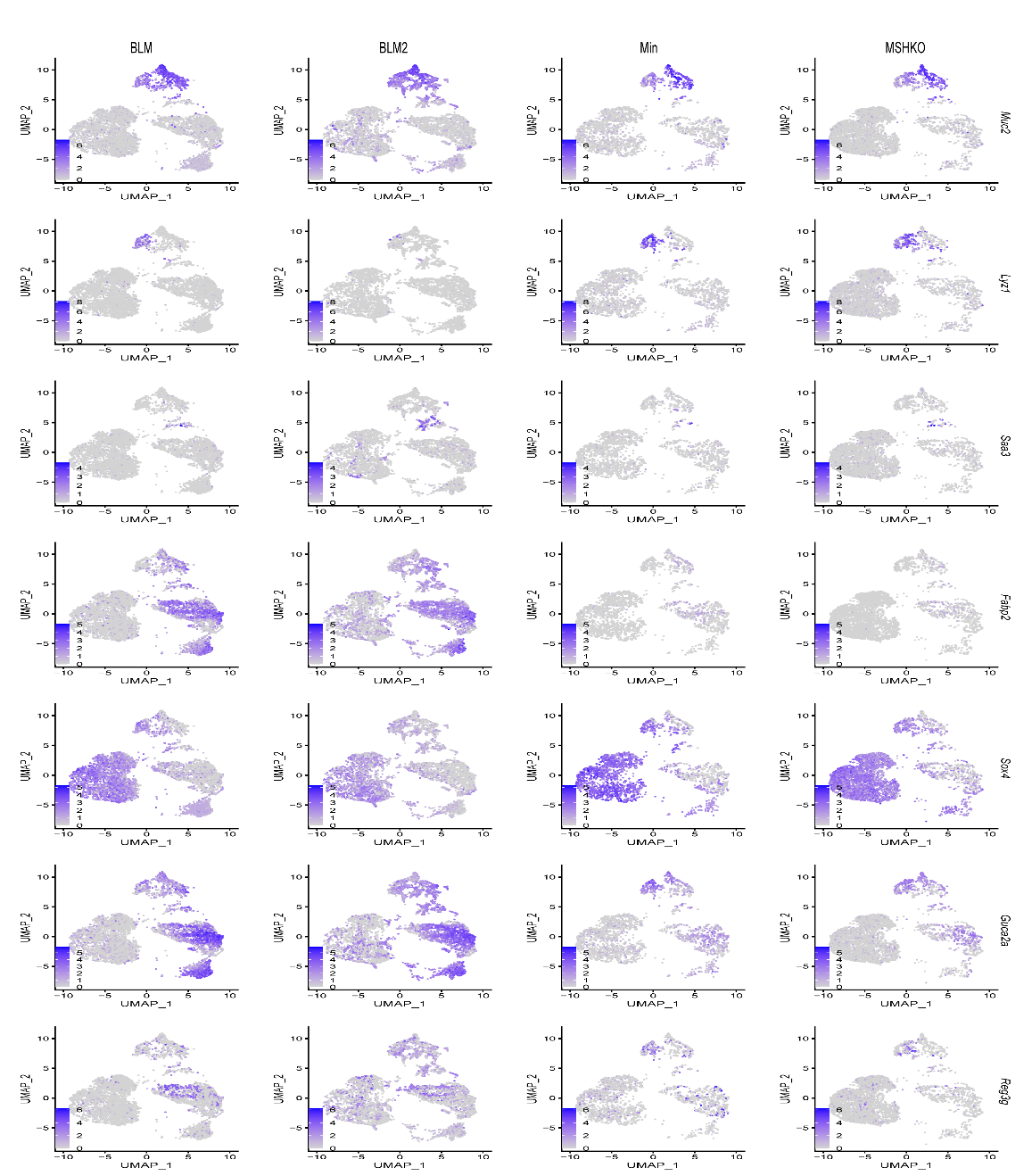

Figure S3. Featureplot of normalized gene expression of different genes in BLM, BLM2, Min, and MSH2KO colon epithelial tumors. Color intensity represents normalized gene expression.

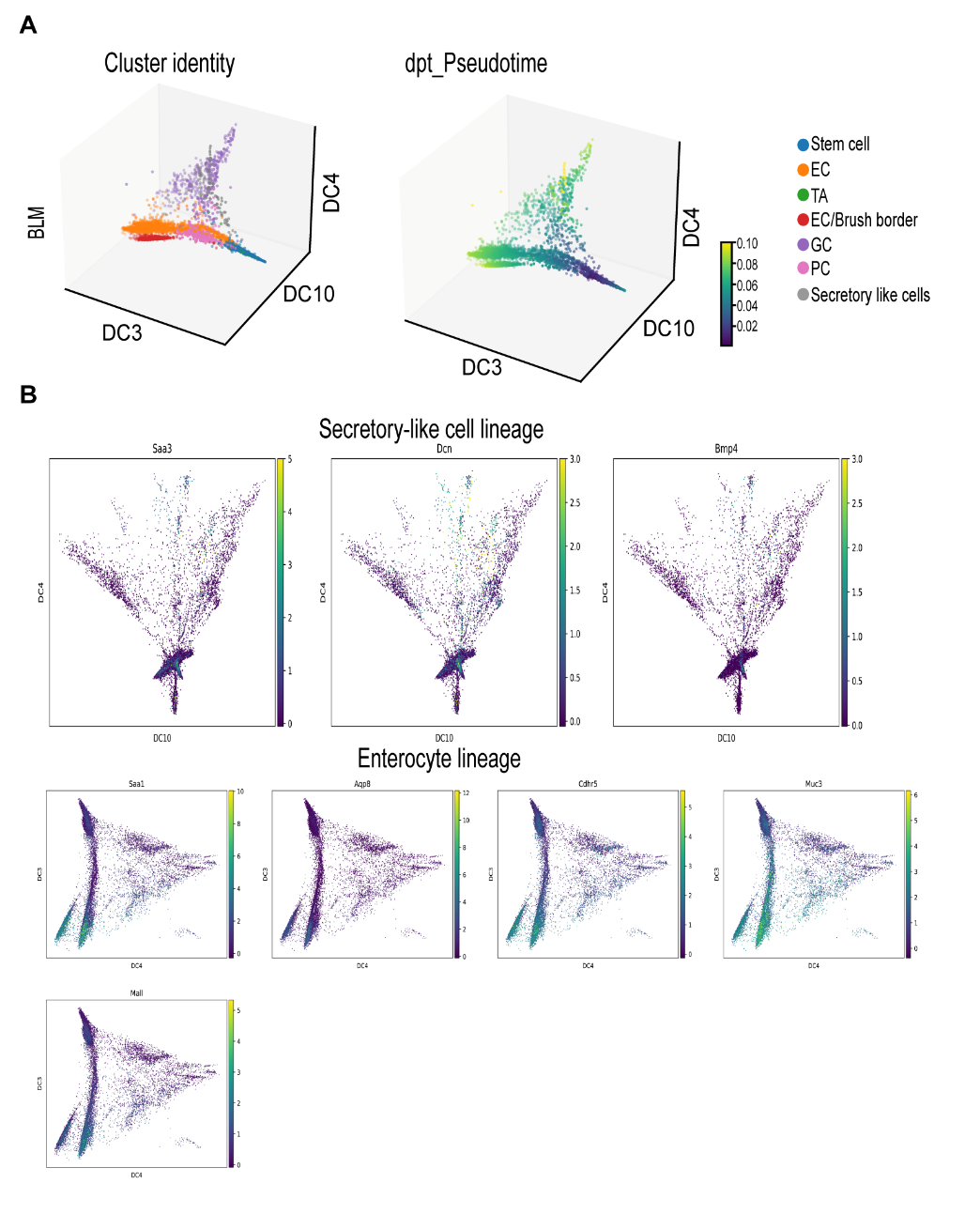

Figure S4. The diffusion map captures the differentiation trajectory of BLM tumors. (A) The diffusion-map embedding of BLM colon epithelial tumor is colored by cell type (left, Cluster identity) and diffusion pseudotime (right, dpt_Pseudotime) (enterocyte cells (EC), transit-amplifying cells (TA), goblet cells (GC), Paneth cells (PC)). The colors represent represent pesudotime value. (B) Expression of regional markers of indicted cell lineages in combined BLM2, BLM, Min, and MSH2KO colon epithelial tumors. Colors represent normalized gene expression.

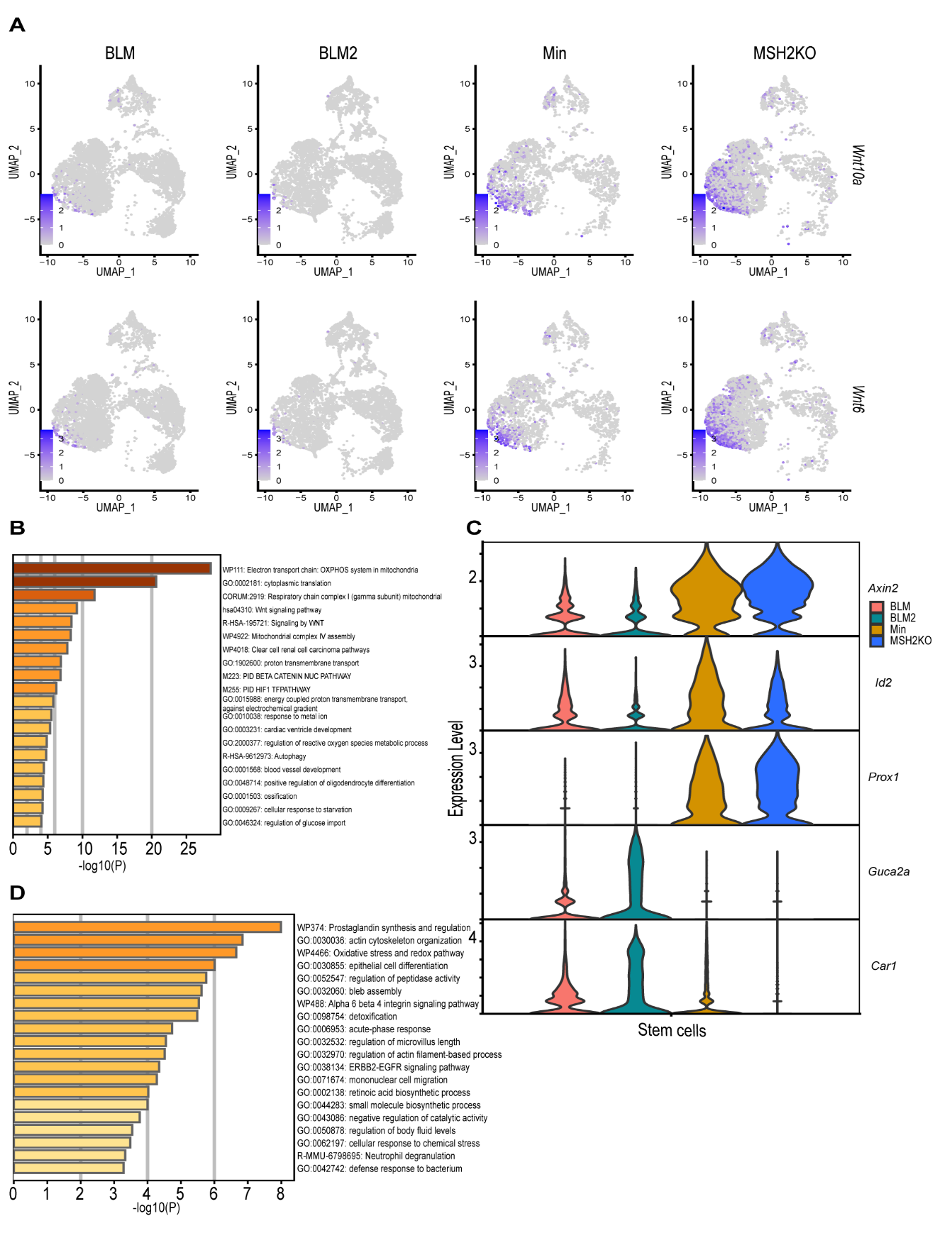

Figure S5. Stem cells of BLM and MSH2KO colon epithelial tumors have low and high WNT signaling activity, respectively. (A) Featureplot of normalized gene expression of *Wnt10a* and *Wnt6* in BLM, BLM2, Min, and MSH2KO colon epithelial tumors. Color intensity represents normalized gene expression. (B) Functional gene annotations for differentially expressed genes (DEGs) in the stem cell populations of MSH2KO versus Min generated by Metascape. (C) Violinplot of normalized gene expression of indicated genes in stem cell clusters of BLM, BLM2, Min, MSH2KO colon epithelial tumors. (D) Functional gene annotations for DEGs in the stem cell populations of BLM2 versus Min generated by Metascape.

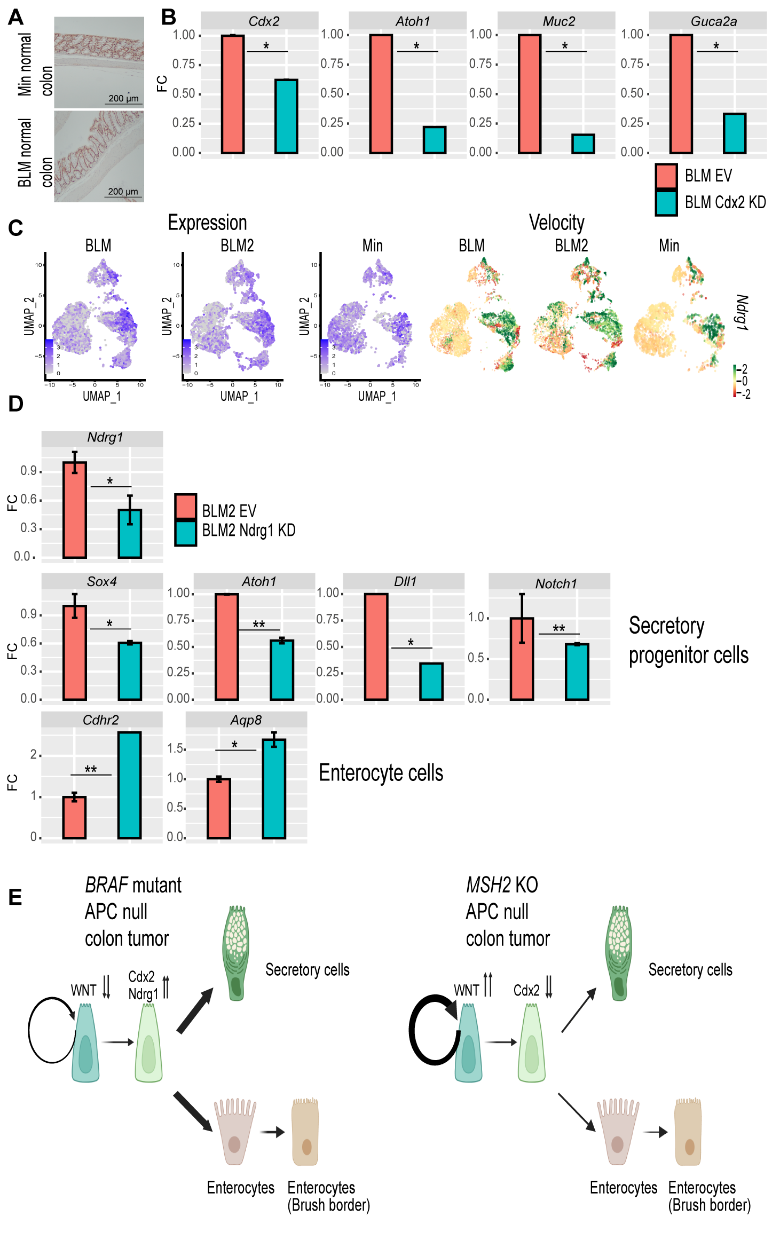

Figure S6. *Cdx2* and *Ndrg1* are involved in BLM colon tumor epithelial cell differentiation. (A) Representative CDX2 IHC in BLM normal colon and Min normal colon (Scale bar, 200 μm). (B) Gene expression of the indicated genes by RT-qPCR in BLM organoids with empty vector (EV) or Cdx2 knockdown (KD). Expression of all the genes was normalized to the housekeeping gene *Ppia* and then to BLM EV organoids. Results are represented as the mean of 3 biological replicates +/− SEM. (C) Featureplot of *Ndrg1* expression (left) and velocity (right) in BLM, BLM2, and Min colon epithelial tumors. Colors represent normalized gene and velocity expression. (D) Gene expression of the indicated genes by RT-qPCR in BLM2 organoids with EV or Ndrg1 knockdown (KD). Expression of all the genes was normalized to the housekeeping gene *Ppia* and then to EV. Results are represented as the mean of 3 biological replicates +/− SEM. (E) The model in which high expression of differentiation-driving transcription factors such as *Cdx2* and *Ndrg1* promotes the differentiation of BLM colon tumors while increased WNT signaling activity increases stem cells in MSH2KO colon tumors. Significance was determined by paired t-test. *p <= 0.05, **p <= 0.01.

| Gene | Taqman assay |
| --- | --- |
| *cdx2* | Mm01212280_m1 |
| *Ndrg1* | [Mm07295892_m1](https://www.thermofisher.com/taqman-gene-expression/product/Mm07295892_m1?CID=&ICID=&subtype=) |
| *Atoh1* | Mm00476035_s1 |
| *Muc2* | Mm01276696_m1 |
| *Dcn* | Mm00514535_m1 |
| *Aqp8* | [Mm00431846_m1](https://www.thermofisher.com/taqman-gene-expression/product/Mm00431846_m1?CID=&ICID=&subtype=) |
| *Notch1* | MM00627185_M1 |
| *Sox4* | Mm00486320_s1 |
| *Dll1* | Mm01279269_m1 |
| *Lgr5* | Mm00438890_m1 |
| *Axin2* | Mm00443610_m1 |
| *Guca2a* | Mm00433863_m1 |
| *Cdhr2* | [Mm01344960_m1](https://www.thermofisher.com/taqman-gene-expression/product/Mm01344960_m1?CID=&ICID=&subtype=) |
| *Ppia* | Mm02342430_g1 |

Table S3. Taqman qPCR assays
